# Patterns of population structure and genetic variation within the Saudi Arabian population

**DOI:** 10.1101/2025.01.10.632500

**Authors:** D.K. Malomane, M.P. Williams, Leqi Tian, Ji Tang, C.D. Huber, S. Mangul, M. Abedalthagafi, C. W. K. Chiang

## Abstract

Both the demographic history and cultural practices influence the pattern of genetic variation, sometimes in opposing manners. For Saudi Arabians, being situated at the hub connecting multiple continents is expected to increase heterogeneity and diversity due to cross-continental migrations. On the other hand, cultural practices such as endogamy to promote social stability may have also promoted regional isolations and reduced heterogeneity. To better understand the genomic impact of these potentially opposing forces, we genotyped and sequenced 3,352 and 302 individuals, respectively, from Saudi Arabians to study their population structure and admixture history, and patterns of genetic variation. We identified twelve genetic sub-clusters that correlated with geographical regions, differentiated by distinct components of ancestry based on comparisons to modern and ancient DNA references. These sub-clusters also showed variation across ranges of the genome covered in runs of homozygosity, reflecting potential differences in kinship or marital practices, as well as differences in population size changes over time. Using 25,488,981 variants found in whole genome sequencing, we found that the Saudi do not show the depletion of rare alleles typically observed in isolated populations, though they do show the expected pattern of enrichment of alleles bioinformatically annotated as deleterious when compared to Africans/African Americans and Non-Finnish Europeans from gnomAD. Saudi sub-clusters with greater inbreeding and lower effective population sizes showed greater enrichment of deleterious alleles as well. Taken together, our results suggest that Saudi’s history and culture impact its pattern of genetic variation and potentially to the population health. We also made available the allele frequency estimates of alleles discovered in our samples so to start a foundation on which to interpret medical- and pharmaco-genomic findings from these populations.

## INTRODUCTION

Saudi Arabia is the largest country in the Arabian Peninsula (AP), the major hub that connects Africa, Asia and Europe. Despite their central location, Saudi Arabians have been relatively under-represented in large-scale genomic studies to understand the genetic architecture of complex traits and diseases. The largest publicly available database for genetic variation in this region is the 2,884 and 147 individuals with summarized exome and whole genome sequencing data, respectively, of “Middle Eastern” origin from gnomAD (Karczewski et al. 2020; Chen et al. 2024) and 161 whole genome sequenced individuals across four countries from the Human Genome Diversity Panel (Bergström et al. 2020)(though also see references (Project Team 2015; Almarri et al. 2020; Mbarek et al. 2022) for other sequencing efforts). Generating the genomic data and characterizing its pattern is the first step towards understanding the genotype-phenotype relationships for these populations.

Both demographic history and cultural practices / human customs can shape the pattern of genetic variation genome-wide. The genetic diversity of today’s Arabians is shaped by a complexity of ancestries from historic split and admixture events. The AP is considered one of the initial sites of historic human migration out of Africa (OOA), with presence of human footprints reported at least since 50 – 60 thousand years ago (kya) and as early as 85 – 120 kya (Armitage et al. 2011; Fernandes et al. 2012; Henn et al. 2012; Rodriguez-Flores et al. 2016; Almarri et al. 2020). The AP is also hypothesized (Lazaridis et al. 2014; Lazaridis et al. 2016; Ferreira et al. 2021) to be one of the most likely homelands of a hypothesized deeply diverged “ghost” ancestry, the Basal Eurasians, who are thought to have diverged from the non-Africans shortly after the OOA. This lineage is then thought to have remained isolated until experiencing a later admixture in the Middle East around 38kya (Vallini et al. 2024) whereby Arabian and other present-day Middle Eastern populations are believed to have descended in-part from this population and carry a higher proportion of this ancestry (Lazaridis et al. 2016; Ferreira et al. 2021). Given its hub location geographically, Arabians have also experienced series of admixtures, and the present-day Arabians could have shared ancestries with various groups including Africans, South Asians, Europeans, Levantines, and Iranians (Fernandes et al. 2019; Almarri et al. 2020; Martiniano et al. 2024). For these geographical and historical reasons, one may expect Arabian genomes to show greater heterogeneity and diversity.

On the other hand, despite the rich history of ancestries and being in a major geographical hub, for centuries the genetic pool of the Arab countries and the Greater Middle East (GME) have been greatly influenced and refined by mating practices. Arab countries have a high rate of endogamous and consanguineous marriages (Tadmouri et al. 2009; Khayat et al. 2024), especially in Saudi Arabia with rates as high as 58% (el-Hazmi et al. 1995; Ben Halim et al. 2013). These endogamous marriages preserve family structure and strengthen bonds, and can ensure cultural, religious, financial and social stability (Bittles 2008; Ben Halim et al. 2013; Alkuraya 2014). Many of the consanguineous marriages are found between first cousins (e.g. 28.4% (el-Hazmi et al. 1995)), but are also found extended to members of the same or related tribal groups. Endogamy leads to regional genetic isolation and population substructure. A recent study analyzing the population structure of Saudi Arabia showed a signature of tribal stratification within the population (Mineta et al. 2021). Co-inheritance of recessively-acting alleles with functional consequences could increase the prevalence of genetic disorders, some of which have indeed been observed in Saudi Arabia (Aleissa et al. 2022; Temaj et al. 2022). While in the long run these deleterious recessive alleles may be exposed to purifying selection due to increased homozygosity (Hedrick and Garcia-Dorado 2016; Delatycki et al. 2020), consanguinity and/or reproductive compensation have been suggested to counteract the effectiveness of purifying selection in endogamous populations (Overall et al. 2002; Alsalem et al. 2013; Greater Middle East Variome Consortium et al. 2016; Sahoo et al. 2021). Therefore, the long-term isolation could lead to reduced heterogeneity and enrichment of bioinformatically-annotated deleterious alleles (Ober et al. 1999; Castellano et al. 2014; Lohmueller 2014; Simons and Sella 2016). That is, cultural practices may counteract the impacts due to geography and ancestry, resulting in an abundance of deleterious alleles.

Prior genomic studies of Saudi Arabia have described broad population structure and identified signatures of tribal stratification (Mineta et al. 2021), but have not been able to address how that structure relates to the formation of the Saudi gene pool over deep time, nor how it translates into measurable genomic risk. We tackle both questions. By integrating whole-genome sequencing with ancient DNA from Epipaleolithic through Medieval Arabian Peninsula and surrounding populations, we reconstruct the ancestry components and admixture timing that shaped each Saudi sub-cluster, moving the field from pattern description to mechanistic history. By characterizing allelic architecture using three independent functional annotation tools across 25,488,981 variants, we directly quantify the net impact of admixture and endogamy on deleterious variant burden, a question not yet addressed by genomics studies in Saudi Arabians. Finally, by identifying nearly 2.5 million variants absent from gnomAD, 19.63% of which are predicted deleterious, we demonstrate that current reference databases are systematically incomplete for this population in ways that matter clinically. By bridging the gap between deep-time demography and modern functional variation, this study provides a framework for understanding how the intersection of history and culture dictates the landscape of human disease.

## RESULTS

### Genetic substructure of Saudi Arabians

We merged 3,352 genotyped Saudi individuals after quality control (see **Methods**) with 302 whole genome sequencing (WGS) samples and explored the population structure based on 603,833 shared segregating sites. Principal component analysis (PCA) on the combined set revealed substantial structure within our sample population (**Figure S1**), consistent with geographical isolations within the population. We defined discrete sub-population for downstream comparisons and analysis by performing clustering analysis which suggested that twelve genetic sub-clusters within the Saudi population best fit the data (**Methods; Figure S2A**). We found that the 12 clusters corresponded to geographical regions within Saudi Arabia, with each cluster generally consisting of a majority of its members from a single geographical region (Central, West, North, South, or East) whether using harmonized regional labels or self-reported labels when available (**Methods**; **Table S1, Figure 1A** and **Figure S2B**). The main exceptions are clusters 11 and 12, both of which consisted of individuals from multiple prevalent regions. In regard to their unique genetic structure, cluster5 from the Western region appeared to be most differentiated from the rest of the cohort (**Figure 1A**; also see PCs 6 and 7 in **Figure S1**). Multiple genetic clusters can be affiliated to the same geographic regions (e.g. clusters 2, 3, and 9 from Central region; 4, 5, 7, and 10 from the Western region, etc.), reflecting the limited demographic resolution of our data, since we do not have access to specific tribal affiliation of each participant due to privacy protections. Even though multiple tribes can inhabit a specific region and inter-tribal marriages within a region is expected to be limited (Mineta et al. 2021), our definition of genetic clusters may not necessarily correspond to distinct or closely related tribes. Instead, they are used to reveal broad geographical and genetic patterns in our study.

**Figure 1.**
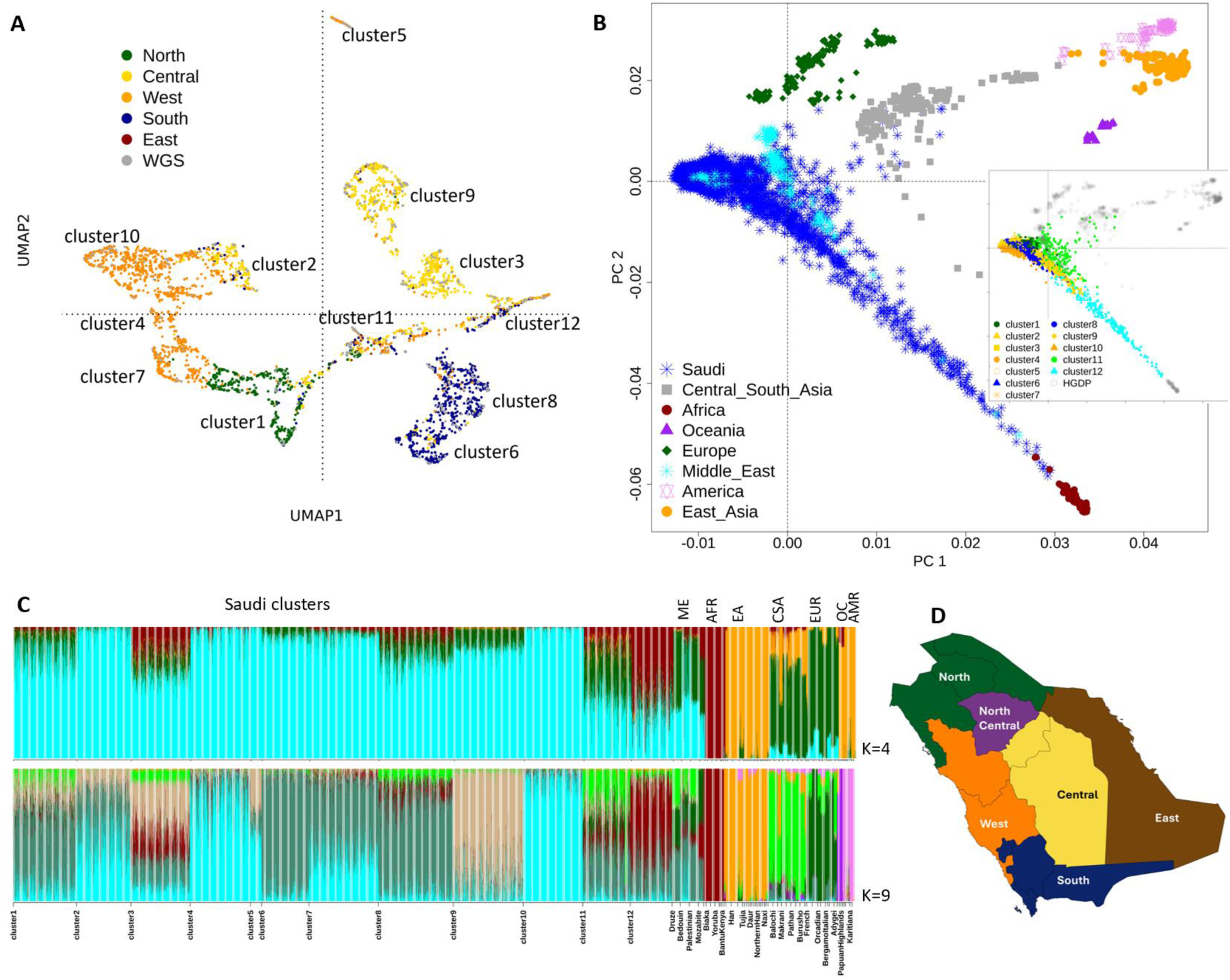
The genetic structure of Saudi Arabians and its relation to global populations. (A) A two-dimensional UMAP of Saudi Arabians based on the top 10 principal components. Each individual is colored based on the affiliated tribal region (see (D)). WGS samples did not have self-reported or harmonized tribal affiliation and are assigned their own color. (B) PCA of Saudi Arabian clusters and HGDP populations. Saudi Arabians are grouped in a single group. Inset shows clusters 1 - 10 colored according to the most prevalent tribal region represented in the cluster (see **Table S1**). Because clusters 11-12 has no single dominating tribal region, they were assigned distinct separate colors. (C) Admixture analysis of Saudi Arabian clusters and HGDP populations for K = 4 (top) and K = 9 (bottom). ME – Middle Eastern, AFR – African, EA – East Asian, CSA – Central & South Asian, EUR – European, OC – Oceania, AMR – American. The names of Saudi clusters and HGDP populations are shown on the bottom X-axis. However, due to limited space some of the labels for smaller populations from HGDP are omitted. Grouped regional labels are shown on the top X-axis of plots. We show the admixture results of the Saudi clusters alone in **Figure S3B**. (D) A regional map of Saudi Arabia with matching colors to the regional labels in (A) and (B).

### Genetic ancestries that can increase diversity

Situated at the crossroad of the African, Eastern and Western Eurasian continent, one may expect the Saudi Arabians to represent multiple components of ancestries and overall exhibit greater genetic diversity than populations in Europe or Asia. To investigate the genetic and ancestral diversity, we compared the Saudi clusters to the populations from the Human Genome Diversity Panel (HGDP) (Bergström et al. 2020). Consistent with previous reports (Mineta et al. 2021; Elliott et al. 2022), the Saudi individuals clustered between Africans, Central & South Asians and Europeans and were the most distant to East Asians (**Figure 1B**). The Saudi’s genetic affinity towards African reference individuals, implicated by a clinal distribution in the first two PCs, are largely driven by cluster12 and cluster3 (**Figure 1B** inset), while cluster11 showed a mixed affinity towards Europeans, Africans and Central & South Asians. The remaining clusters co-localized mainly with the Middle Eastern reference individuals from the HGDP panel (**Figure 1B**).

Our observation from PCA is also corroborated by unsupervised ADMIXTURE analysis at two different resolutions of K (K = 4 and 9), where clusters 11 and 12 exhibited the highest levels of admixture (**Figure 1C, Figure S3A, S3B,** and **Table S2**). At K = 4 where major continental ancestries relevant to Saudi Arabians are differentiated, the most dominating ancestry in the Saudi clusters was one largely shared with the HGDP Middle Eastern populations (Druze, Bedouin, Mozabite, and Palestinian; cyan ancestry in **Figure 1C**, top). Clusters 12, 11, and 3 had on average less than two-thirds of this Middle Eastern (ME)-like ancestry component and were enriched with African-like (red) and/or European-like (green) ancestries. At K = 9, the relationship between cluster11 and the Central & South Asians (CSA) that we observed on PCA can also be observed (**Figure 1C** bottom, **Table S3**), where cluster11 carried more (average proportion = 0.223) of such CSA-like ancestry compared to other clusters (average proportions less than 0.1).

To further elucidate Saudi genetic history and its ancestral components, we integrated whole genome sequencing data from 302 Saudi individuals with ancient DNA (aDNA) datasets (1240k AADR v62.0 (Mallick et al. 2024); **Methods**). Despite its importance, the Arabian Peninsula remains underrepresented in aDNA records with available samples limited to two locations: Bahrain in the Persian Gulf (four Tylos-period individuals, ∼2200–1400 BP; (Martiniano et al. 2024)) and Medieval Soqotra at the mouth of the Gulf of Aden (twelve individuals, ∼1300 BP; (Sirak et al. 2024)). Using D-statistic symmetry tests of cladality (**Methods**), we assessed whether each of the 12 Saudi sub-clusters are consistent with a model of population continuity with these ancient Arabian groups (**Figure S4, Table S4**). Across all tests, we rejected a model of strict population continuity between ancient Arabian groups and present-day Saudi populations, finding instead a consistent pattern of African and Levantine-related ancestry input. This observation was corroborated through affinity *D*-statistics which showed that clusters 1–10 have increased genetic affinity to ancient Levantine populations (e.g., Syria_TellQarassa_Umayyad.SG, Israel_Natufian.AG, and Neolithic/EBA samples; **Figure S5, Table S5**) dating to the 7th and early 8th centuries, Epipaleolithic, and Neolithic to Early Bronze Age periods (∼1200 to 11,500 years ago). Moreover, outgroup *f*3-statistics revealed all clusters, with the exception of cluster 12, showed maximal shared drift along a Levant-Anatolia-Mediterranean-Balkan axis (**Figure 2A**, **Figure S6, Table S6**). These results suggest that since the Late Tylos period, the ancient Arabian gene pool received additional input from a population best represented by Neolithic or Early Bronze Age Levantine samples.

**Figure 2:**
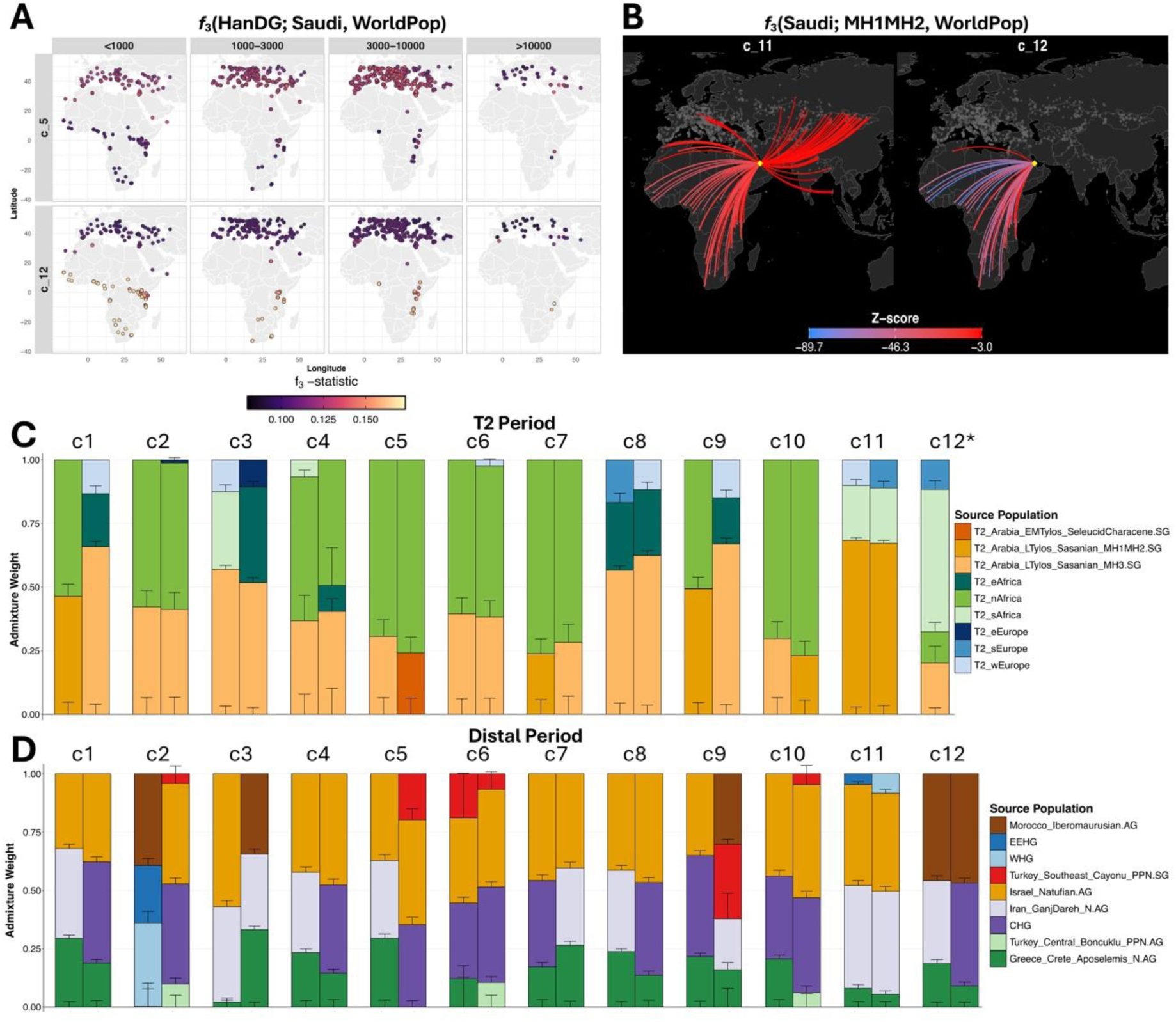
Genetic affinities and admixture modeling of Saudi clusters. (A) Outgroup f3-statistics of the form f3(Han.DG; Saudi, WorldPop) mapping shared genetic drift between specific Saudi clusters c_5 and c_12 and global populations, stratified by temporal periods (years BP). Darker colors indicate higher shared drift. See **Figure S6** for full results across Saudi clusters. (B) Admixture signals illustrated via admixture f3-statistics, demonstrating gene flow trajectories and affinities across the region. The color scale denotes the Z-score, with red lines indicating significant admixture. Ancient Bahrain MH1MH2 samples were used in admixture f3-statistics here; for admixture f3-statistics involving other ancient Arabian samples and Saudi clusters, see **Figure S7**. (C, D) Genetic ancestry modeling using *qpAdm*, displaying the admixture weights based on more proximal (T2 in C) or deep (Distal in D) temporal source populations. Error bars denote +/- 1 standard error. Top two plausible models ranked by p-value for each Saudi cluster, if available, are shown from left to right.

On top of the Levantine-related West Eurasian ancestry, several Saudi clusters exhibit a major sub-Saharan African ancestry component, represented in the ancient DNA dataset by Eastern and Northeastern African populations, including the Kenya-associated pastoralists, foragers, and their modern proxies (e.g. Kenya_KisimaFarmA5_PN, Kenya_HyraxHill_PN, Kenya_Kakapel_LIA, Kenya_Somali) as well as populations from the coastal / Great Lakes region (e.g. Tanzania_Lindi_Swahili, Tanzania_Pemba_600BP/1400BP, Uganda_Munsa_LIA) (**Figure S4**). This pattern is most pronounced in cluster 12, followed by cluster 3. Outgroup *f*3-statistics identify cluster 12 as a clear outlier in the estimated shared drift with ancient sub-Saharan African samples over at least the last 10,000 years (**Figure 2A**), and formal admixture *f*3-statistics confirm the signal of a mixed ancient Arabian + sub-Saharan African ancestry in clusters 3, 8, 11, and 12 (**Figure 2B, Figure S7, Table S7**). Taken together, the shared drift and admixture tests with ancient samples confirmed the strong affinity between some Saudi sub-clusters and sub-Saharan Africans inferred from modern reference populations described above (**Figure 1C**).

Other than sub-Saharan African ancestry, cluster 11 exhibits an additional distinct pattern. Its affinity D-statistics profile shifts away from an exclusively Levantine-related ancestry toward a mixed Anatolian and European Neolithic/Mediterranean and CSA profile (**Figure S5**). This is most evident in the admixture *f*3-statistic, which reveal exclusive connections to Eastern European, CSA, and Steppe groups (**Figure 2B, Figure S7**). These findings corroborate the European and CSA-like ancestry profile observed in ADMIXTURE analysis with modern reference samples described above (**Figure 1C**).

We employed qpAdm to model the genetic ancestry of modern Saudi Arabians through a time-stratified lens whereby we grouped the available aDNA reference populations into thirteen broad geographical regions encompassing major regions in Europe, Caucasus, Levant, Anatolia, Arabia, and Africa (**Methods, Tables S8-S11**). Our analysis successfully modeled the genetic ancestry profile of the Saudi Arabians at two different temporal strata (T2: 1,000-3,000 BP; Distal: ancestrally representative populations ∼ > 7,500 BP). In the proximal period to the present (T2; 1,000–3,000 BP), parsimonious genetic models for the Saudi clusters fall into two broad patterns. Seven clusters (c1, c2, c5, c6, c7, c9, c10) fit a simple two-source structure of ∼24–49% Arabian-related ancestry and ∼51–76% North African-related ancestry (**Figure 2C**). Within these models, the Late Tylos groups MH1-MH2 and MH3 appear most frequently as the Arabian source, with the Early-to-Middle Tylos sample observed less frequently. These results are consistent with affinity D-statistics (**Figure S5**), where the North African aDNA sources appear to capture the Levantine-Anatolian and sub-Saharan components. For the remaining clusters, an additional ancestry component from Southern/Eastern Africa (c3, c4 and c11) and/or Europe (c3 and c11) was required to achieve a parsimonious fit (**Figure 2C**). Cluster12 initially failed to fit any tested qpAdm model with up to six sources and required a focused analysis (**Methods**) to identify a single plausible four-source model: Late Tylos MH3 (∼20%), Northern Africa (∼12%), Southern Africa (∼56%), and Southern Europe (∼12%). These results underscore the ancestral diversity and heterogeneity of cluster12, particularly its significant Sub-Saharan ancestry. Such complexity possibly derives from the fact that both cluster11 and cluster12 include individuals from multiple distinct geographical regions.

For the most distant temporal stratum utilizing representative ancestral populations of available regions, a common baseline ancestry structure is shared among many clusters (c1, c3, c4, c5, c7, c8, c9, c10). This baseline is adequately modeled as a three-source mixture: Iranian Neolithic/Caucasus Hunter-Gatherer-related ancestry (up to ∼33%), Neolithic Anatolian Farmer via Greece and Crete-related ancestry (up to ∼30%), and Epipaleolithic Levantine ancestry via Israel Natufian (∼32-57%) (**Figure 2D**). These findings are consistent with the demographic modeling of Marchi et al. (Marchi et al. 2022), wherein three meta-populations emerged following the Late Glacial Maximum (LGM) recolonization of the Middle East. Cluster12 departs from the three-source ancestry baseline by integrating a Morocco-Iberomaurusian-related component (∼46%) with Neolithic Anatolian (∼19%) and Iranian/Caucasus (∼35%) elements. This Iberomaurusian-related ancestry, which possesses distant genetic connectivity to Levantine Epipaleolithic Natufians, appears to also capture ancestry related to the hypothesized ‘Basal Eurasian’ lineage – a ghost lineage that diverged from the primary out-of-Africa lineage prior to Neanderthal introgression (Lazaridis et al. 2014; Fregel et al. 2018; Van De Loosdrecht et al. 2018; Yang and Fu 2018).

The ancestry of the three remaining Saudi clusters (c2, c6, c11) could not be explained by the baseline structure alone. Cluster11 required a four-source model, adding a European Hunter-Gatherer ancestry component – either Eastern (∼4%) or Western (∼8%) – to the three-source baseline **(Figure 2D**), a finding supported by admixture *f*3-statistics (**Figure 2B; Figure S7**). Conversely, clusters 2 and 6 incorporate varying proportions of a Pre-Pottery Neolithic Mesopotamian source (c2 = < 1%; c6 = ∼19%), which itself represents a composite of Levantine, Anatolian, and CHG/Zagros lineages (Altınışık et al. 2022). The observation with clusters 2 and 6 suggests that while the currently available distal sources best represent the broad ancestral diversity of the Saudi clusters during the Neolithic-Paleolithic periods, they are likely still imperfect proxies.

Finally, previous research has identified genetic component among present-day AP populations related to Basal Eurasian ancestry (Fregel et al. 2018; Van De Loosdrecht et al. 2018). Using *f*4-statistics (**Methods**), we estimated the relative amount of Basal Eurasian ancestry using various ancient and present-day African outgroups and identified a latitudinal gradient where Basal Eurasian proportions decrease in the northern Middle East, placing most Saudi clusters alongside the southern Levant and Arabian Bedouins (**Figure S8A, Table S12**). At the extreme of this gradient is cluster12 who possesses estimates comparable to present-day East Africans (e.g., from Eritrea and Ethiopia, and the Afar and Shaigi peoples). Whilst this outlier effect is more pronounced when using a present-day African source as the outgroup (e.g., Yoruba) rather than the ancient North African Basal Eurasian surrogate, Iberomaurusian (**Figure S8B, Table S12**), distinguishing African admixture from putative Basal Eurasian ancestry remains a challenge.

### The social structure that can reduce genetic variation

Genetic ancestries, preeminently driven by genetic similarity to African-related ancestries but also possibly by Basal Eurasian ancestries, could dictate the pattern of genetic variation of the Saudi Arabians. On the other hand, another major force not to be overlooked is the cultural practices, such as endogamy, that establishes the social structure in this population and could impact the pattern of variation. We thus examined one of the genetic hallmarks of endogamy, runs of homozygosity (ROH). Comparing the number of ROH (NROH) vs. the sum total length of ROH (SROH), we found that the Saudi Arabians exhibit a diverse distribution of ROH between clusters and between the individuals (**Figure 3A** and **3B**). Across Saudi clusters, the median SROH ranged from 38.12 Mb to 232.6 Mb, while the median NROH ranged from 42 to 150 ROHs. Clusters 12, 11, and 3 had the shortest mean length and smallest mean number of ROHs (**Figure 3A**), consistent with greater admixture from more diverse African ancestral populations (**Figure 1C**). Cluster5 had the highest burden of ROH, with highest average number and total length of ROH, reflecting a consequence of both long-term small effective population size and/or consanguinity (Ceballos et al. 2018) (**Figure 3A**).

**Figure 3.**
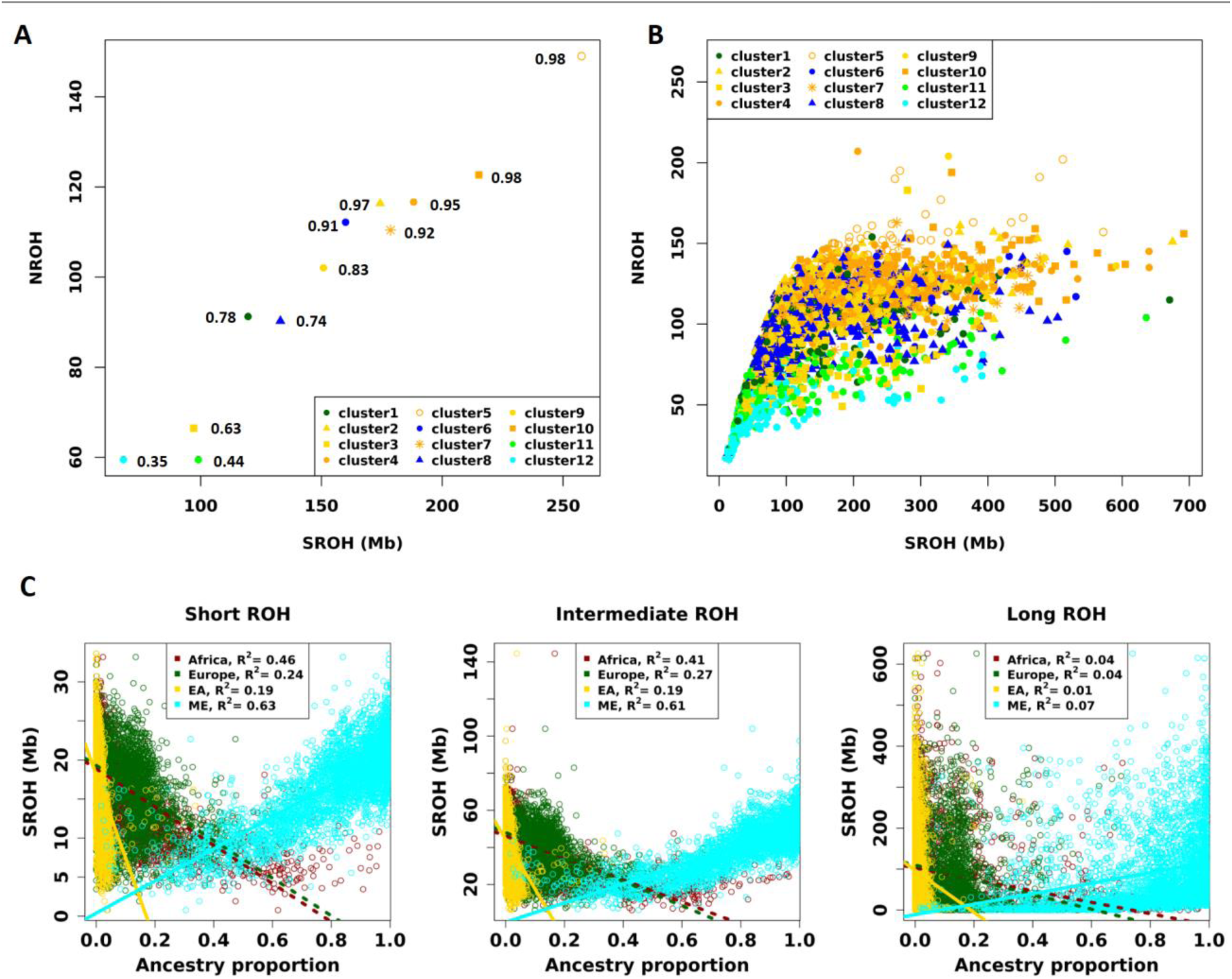
Runs of homozygosity in Saudi Arabians. (A) Average total length and number of ROH per cluster. The numbers next to the symbol represents the mean ME-like ancestry proportion. (B) Total length and number of ROH per individual across the Saudi Arabian cohort. For (A) and (B), symbols are colored by the geographical region associated with each cluster (Figure 1D). (C) Total length of ROH vs ancestry proportion per individual stratified by three length classes of ROHs. Short ROHs indicate homozygosity from ancient or distant ancestry, i.e. background relatedness. Intermediate ROHs likely arise from background relatedness with moderate level of inbreeding from past few generations, often due to reduced population sizes or reproductive isolation (e.g. due to geographic or cultural preferences), or from recent bottlenecks followed by recovery. Long ROHs arise through recent inbreeding and are common in populations with high levels of consanguinity (Pemberton et al. 2012; Thompson 2013; Ceballos et al. 2018). ROH – Runs of homozygosity, ME - Middle Eastern, EA – East Asia.

We also followed a previous approach (Pemberton et al. 2012) and divided the ROHs based on length into short, intermediate, and long classes. When classified by the sizes, we can observe that the overall pattern of NROH vs. SROH (**Figure 3B**) are driven by the long ROHs (**Figure S9**), where contributions of SROHs are driven by fewer but longer ROHs in the long length class. Long ROHs tend to arise from recent inbreeding. The pattern of NROH and SROH across Saudi sub-clusters are also consistent for ROH of the short and intermediate length classes (**Figure S10A** and **S10B**), but varied for the long ROH class, implying a different pattern of recent consanguinity across sub-clusters in contrast to their shared ancient demographic events. Therefore, the consequences of consanguinity in not only increasing SROH but also increasing the variance of SROH in a population (Ceballos et al. 2018; Ceballos et al. 2021).

When examined in light of estimated genetic ancestry (at the continental level, K = 4; **Figure 1C**), the pattern of NROH vs. SROH showed a distinct relationship with the proportion of ME-like ancestries, with greater ME-like ancestry also showing greater ROH in length and number (**Figure 3A**). In fact, we found that SROH is positively correlated with ME-like ancestry proportions, and negatively correlated with the proportion of African, European and East Asian ancestries (**Figure S10C**). This observation is seen across length classes of ROHs, though more attenuated for long ROHs (**Figure 3C**). We reasoned that this ancestry effect is again likely reflecting the commonly practiced endogamous marriages and recent consanguinity associated with the ME-like ancestry.

Isolation by endogamy may also be reflected in persistent small population sizes. We leveraged the dense marker information from the 302 WGS individuals to reconstruct genome-wide genealogies and infer the population size trajectories within the Saudi sub-clusters (**Figure 4**). We found that all clusters experienced similar history through the out-of-Africa bottleneck (∼100 kya), followed by a recovery to a local maximum in effective population size (Ne) 10-20 kya, a period consistent with the early Holocene Wet Phase / Holocene Humid Period, characterized by wet conditions which resulted in expansion of lakes and rivers and extensive grasslands (Petraglia et al. 2020). Following the Holocene period, sub-clusters began to diverge around 6-10kya, coinciding with the Arabian aridification, which is responsible for the desert conditions in most of the Arabia as we know it today (Petraglia et al. 2020; Martiniano et al. 2024). Clusters with less ME-like ancestries and stronger signature of admixture (such as clusters 12, 11, 8, & 3), showed less severe decline in Ne compared to those that have high ME-like ancestry. Cluster5 in particular, showed the most severe bottleneck and remained low in Ne in the recent times. Cluster5 appears to resemble the pattern of the tribe labelled as T25 in a previous study (Mineta et al. 2021); both originated from the Western region showing the highest level of inbreeding within the respective study. T25 is said to have been subjected to strict intratribal marriages. Such social practices can indeed result in persistent small Ne as observed here, as well as our observed pattern in ROH (**Figure 3**).

**Figure 4.**
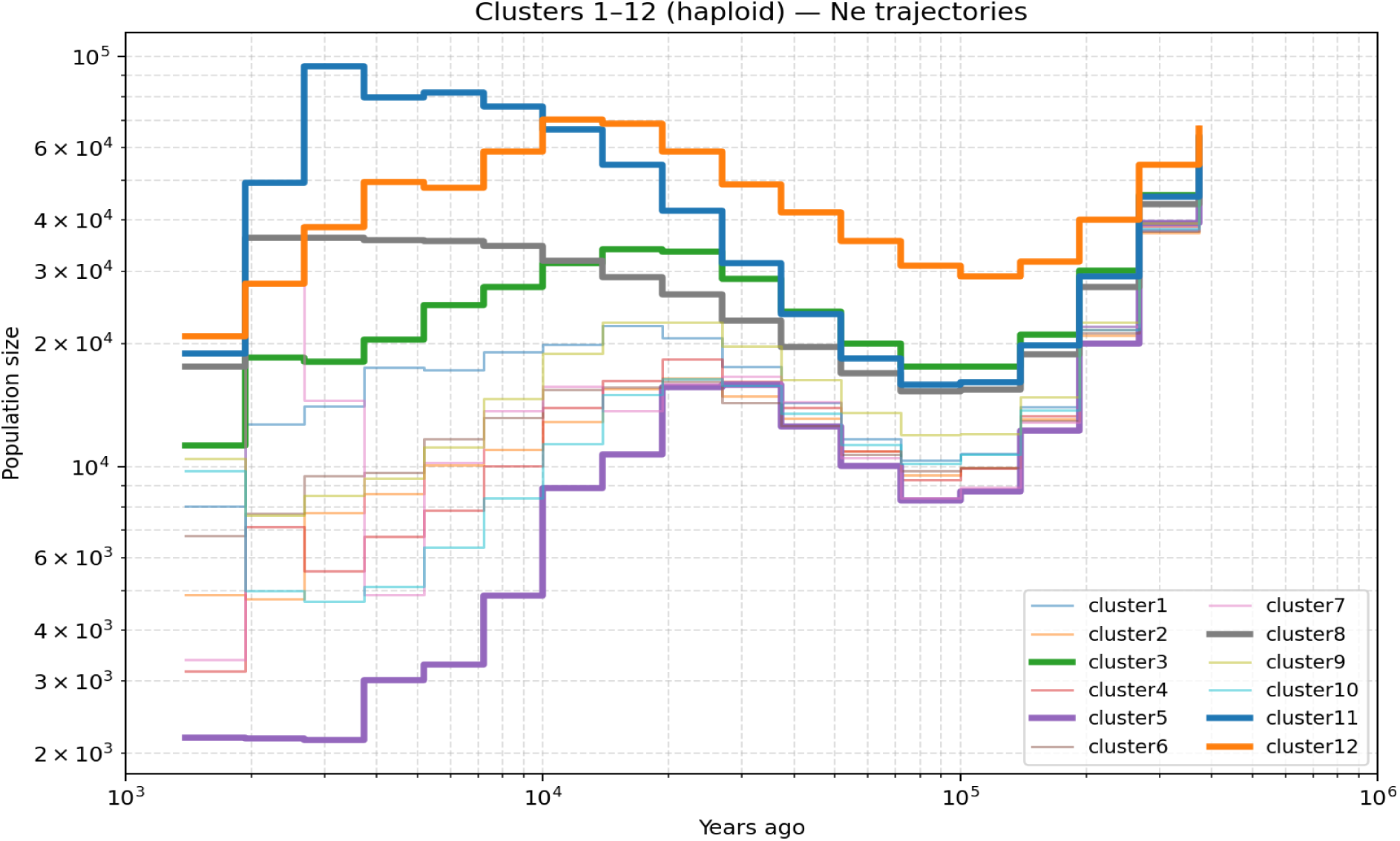
Population size trajectories between the Saudi Arabian sub-clusters. Effective population sizes were computed from genealogical trees using RELATE (see **Methods**). The number of samples per cluster used for the estimates can be found in **Table S1**.

### Allelic architecture of Saudi Arabians

#### Genetic variation in Saudi Arabian WGS data

Having investigated extensively the genetic signature of ancestry, admixture history, and endogamy in the Saudi, we next sought to determine how these demographic and cultural factors influence the patterns of genetic variation in Saudi. There are potentially two competing effects: first, significant contributions from African-related and Basal Eurasian ancestries, at least in some sub-clusters, would increase genetic diversity and facilitate more effective purifying selection. Conversely, culturally embedded practice of endogamy is expected to decrease genetic diversity, reduce effective population sizes, and relax the effect of purifying selection, potentially leading to the enrichment of functionally deleterious alleles. There is empirical evidence of observing an excess of functionally deleterious alleles in isolated populations (Lim et al. 2014; Pedersen et al. 2017; Locke et al. 2019). Driven by bottlenecks, these populations exhibit two hallmark signatures of their allelic architecture: the paucity of rare alleles and the enrichment of deleterious alleles at intermediate frequencies. The pattern has not been examined among populations like the Saudi Arabians, where both admixtures and the long-standing practice of endogamy could both affect the allelic architecture. Here we leverage the whole genome sequences (WGS) from 302 Saudi individuals to investigate the net impact of these opposing forces on pattern of variation.

In total, 25,488,981 autosomal variants were called and retained after quality control (QC) (**Methods**). We first compared allelic frequency spectra and allelic homozygosity in the Saudi Arabians (all WGS individuals) to the Middle Eastern population in gnomAD (gnomAD-MID). The two have relatively similar patterns in the genome-wide alternative allele frequency spectra though Saudi had proportionally slightly fewer common variants (**Figure S11A**). The allele frequencies are highly concordant (r = 0.98) between the two populations (**Figure S11B**), but Saudi Arabians have approximately 2x more homozygous genotypes than gnomAD-MID (e.g. an average of 20% and 10% of the genotypes are homozygous for variants with alternative allele frequency > 5% in Saudi and gnomAD-MID, respectively; **Figure S11A** and **S11C**). The higher proportion of homozygous variants confirms that the Saudi and the gnomAD-MID population are not reflective of the same underlying populations. However, because the frequency spectra and correlation of allele frequencies are highly similar (**Figure S11**), we thus utilize both samples to compare the pattern of variation with gnomAD African/African Americans (gnomAD-AFR) and non-Finnish Europeans (gnomAD-EUR) to better understand the impact of the unique history in the Arabian Peninsula on its current pattern of variation.

#### Distribution of functionally deleterious variants

We annotated the variants using three different annotation tools: VEP (v.110) (McLaren et al. 2016), AlphaMissense (Cheng et al. 2023), and Genomic Pre-trained Network (GPN) (Benegas et al. 2023). AlphaMissense predicts the pathogenicity of missense variants while GPN predicts the deleteriousness for both coding and non-coding variants. The distribution of the variants by functional classes are shown in **Table S13**. Of the called variants, 2,459,950 (9.7%) variants were not previously identified in gnomAD v4.1 (Karczewski et al. 2020; Chen et al. 2024) and thus are potentially novel or Saudi-specific. We refer to these variants as the “previously unknown variants”, or PUVs. As expected, the PUVs are highly enriched with rare alleles (*e.g.* 83% of them are singletons in our dataset, compared to 32% singletons among the known variants; **Figure S12**). In addition, proportionally more PUVs (19.63%) were annotated to be deleterious than known ones found in gnomAD (7.2%). This Implies that the PUVs are not just sequencing errors distributed randomly across the genome, but are enriched for rare variants of functional consequences that are maintained in the Saudi population.

We then compared the allelic architecture of functionally deleterious alleles in the Saudi population to other continental populations from gnomAD. Compared to gnomAD-AFR individuals, the Saudi tend to show proportionally more deleterious alleles than those annotated to be benign or neutral across algorithms (**Figure 5A**), particularly for variants up to ∼5% frequency. Overall, relative to gnomAD-AFR, between the 0.5 - 5% frequency, we found a 13% proportional increase of deleterious (likely pathogenic) alleles annotated by AlphaMissense in the Saudi Arabians compared to 7% proportional decrease of the benign alleles (*P* < 0.01; **Figure 5A**). When annotated by VEP and GPN, at the same frequency range, we observed a consistent pattern i.e. a 3% proportional increase in loss-of-function variants in the Saudi Arabians compared to 10% proportional decrease in neutral (synonymous) ones (*P* < 0.01) by VEP, and an 11% proportional increase in the first percentile of alleles by deleteriousness compared to 3% proportional decrease in the 99^th^ percentile (e.g. the most likely neutral) of alleles when annotated by GPN (**Figure 5A**). We also qualitatively replicated these patterns by comparing the exome samples from gnomAD-MID to gnomAD-AFR (**Figure S13**), or by comparing Saudi to gnomAD-EUR (**Figure S14**).

**Figure 5.**
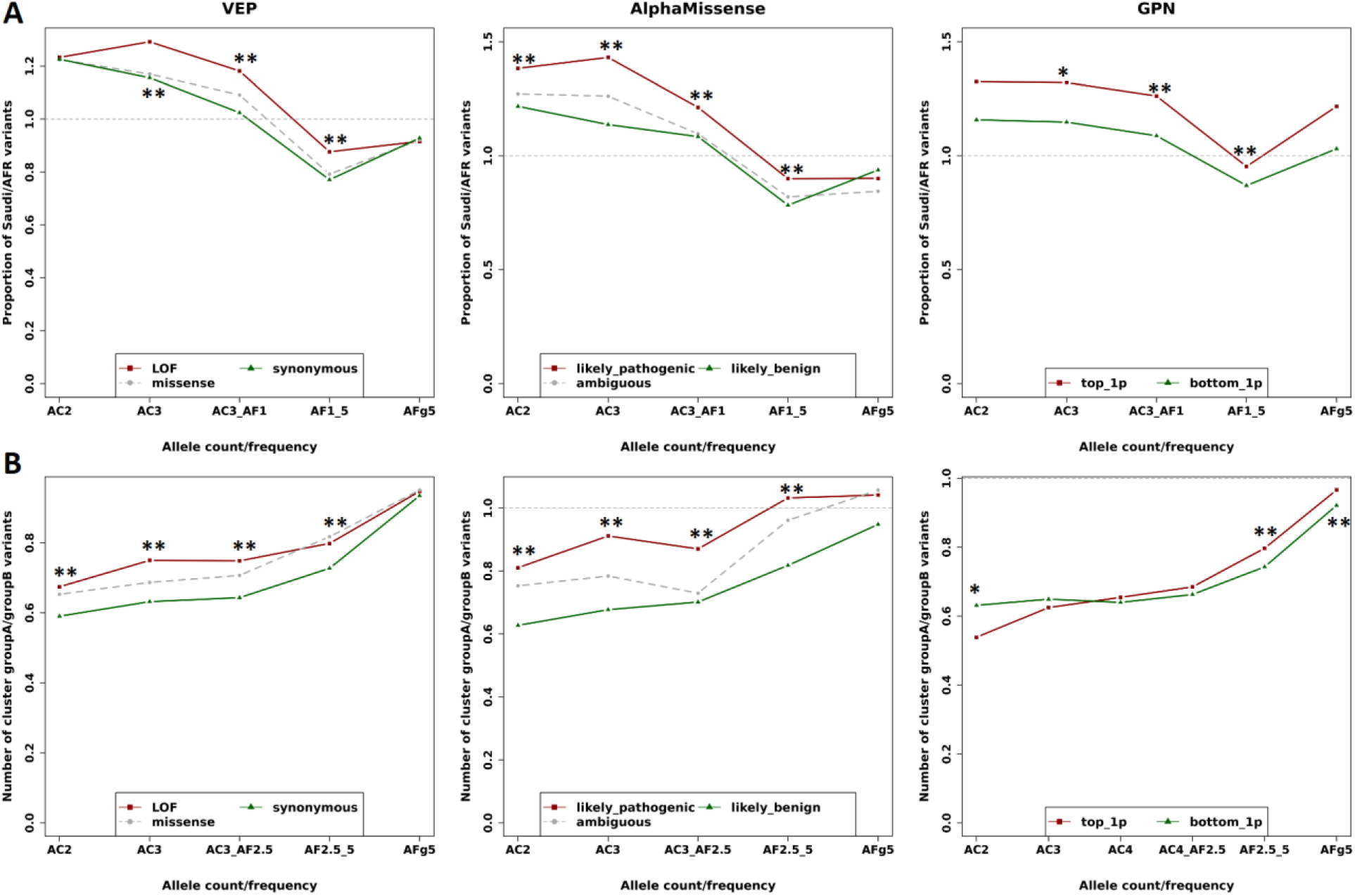
Distribution of minor allele frequency across functional classes. (A) Ratio of Saudi to gnomAD-AFR variants. The sample size of gnomAD-AFR is based on downsampling to Saudi sample size, n = 302. (B) Ratio of Saudi cluster groupA to cluster groupB variants. The sample size of cluster groupB is based on downsampling to groupA sample size, n = 124. Variant functional consequences were annotated based on VEP (loss-of-function, missense, or synonymous variants), AlphaMissense (likely pathogenic, likely benign, and ambiguous), and GPN. AC and AF refer to allele count and allele frequency, respectively. AFg5 refers to allele frequency greater than 5%. Top_1p refers to variants with the top 1% of GPN scores (more deleterious) and Bottom_1p refers to variants with the bottom 1% of GPN scores (more neutral). AFR denotes the gnomAD-AFR sample. LOF refers to Loss of function. ** and * denote frequency bins with significant difference between the most deleterious (red) and most neutral (green) through bootstrapping at *p* < 0.01 and < 0.05, respectively.

We also compared the enrichment of deleterious alleles between Saudi sub-clusters. Because of the smaller number of individuals within each cluster having WGS data (**Table S1**), we grouped the clusters into two groups: groupA which contained clusters with greater amount of SROH and lower effective population sizes (clusters 2, 4, 5, 6, 9, and 10), and groupB which showed less SROH and higher effective population sizes (clusters 12, 11, 3, and 8). We left out cluster1 from this analysis as it tends to fall in the middle of the two groups. GroupA had generally fewer number of variants compared to groupB (**Figure S15**), consistent with its lower genetic diversity, smaller effective population sizes, and greater SROH, and also showed greater enrichment of deleterious alleles (**Figure 5B**).

Notably, across these analyses and particularly when comparing Saudi or gnomAD-MID to gnomAD-AFR or gnomAD-EUR, we did not observe one of the hallmark signatures often seen in other isolated populations – the paucity of rare alleles. In general, both the Saudi and gnomAD-MID had more rare alleles (*e.g.* variants with alternative allele counts of 2 or 3) than the gnomAD-AFR population, potentially due to the admixture history in the Arabian Peninsula, such as through the Basal Eurasian ancestry. Consistent with this hypothesis, we found a correlation between measures of Basal Eurasian ancestry (based on the f4-ratio, **Methods**) and observed heterozygosity per individual (R^2^ = 0.612; **Figure S16A**). However, a similar trend is also observed between estimated African-like ancestries (based on K = 4, **Figure 1C**) and heterozygosity (R^2^ = 0.814; **Figure S16B**). Both a linear mixed-model and nested model comparison testing the association of Basal Eurasian ancestry and heterozygosity were not significant after accounting for African-like ancestries (LMM *P* = 0.519, LRT *P* = 0.5; **Figure S16C-E**). However, together, these ancestry predictors explained a large proportion of the variance in heterozygosity across the dataset (Marginal R^2^ ∼80.5%). This explained variance was almost entirely driven by the African-like ancestry component (Partial R^2^ = 0.698), a finding consistently observed even if we iteratively left one Saudi cluster out of the analysis at a time (**Figure S17**). Importantly, while Basal Eurasian ancestries and African-like ancestries estimated here moderately colinear (variance inflation factor (VIF) = 3.62), it remains unclear whether the excess of rare alleles in Saudi compared to gnomAD-AFR populations is driven by elevated Basal Eurasian ancestry or differences between Eastern- and Western-African-like ancestries.

## DISCUSSION

Scholars have long emphasized the complex demographic histories shaping the genetic architecture of populations across the Arabian Peninsula and have called for improved characterization to better understand regional genetics and health implications (Charati 2021; Elliott et al. 2022). Situated at the crossroads of Africa and Eurasia, Arabian populations are expected to exhibit elevated heterozygosity due to extensive intercontinental interactions. Conversely, deeply rooted traditions of endogamy in Saudi Arabia promote homozygosity, with potential health consequences (Sahoo et al. 2021; Aleissa et al. 2022). Here, we characterize the fine-scale population structure of Saudi Arabia using 3,252 genotyped and 302 whole-genome sequenced individuals sampled across the country, examining how genomic history and social structure jointly shape genetic variation and disease risk.

Integration of our WGS data with published ancient genomes clarifies the formation of the contemporary Saudi gene pool. For most clusters (c1, c2, c5, c6, c7, c9, c10), we identified a recent deviation from ancient Arabian ancestry driven by the integration of Levantine/Anatolian-related components. This shift is reflected in both D-statistics (**Figure S4, Figure S5**) and T2-period qpAdm modeling (**Figure 2C**), where the most parsimonious framework for many clusters involves a two-way mixture of T2_nAfrica (∼51–76%) and ancient Arabian groups (Late Tylos MH1, MH2, or MH3 ∼24–49%). The efficacy of North African samples during this period as a proxy is explained by its comprising of ancient Egyptian (Schuenemann et al. 2017) and Kulubnarti Nubian (Sirak et al. 2021) samples, both of whom harbor significant shared ancestry with Levantine Near Eastern populations. In the case of the Kulubnarti Nubians, this is thought to reflect continuous gene flow along the Nile corridor from 490 BCE to 850 CE.

Superimposed on this base model, African-related admixture distinguishes cluster12 and, to a lesser extent, clusters 3, 8, and 11 from other Saudi clusters. Previous studies attributed African ancestry in Arabia primarily to Bantu-speaking sources from Eastern or Southern Africa, dating to 400–1754 years ago (Hellenthal et al. 2014; Fernandes et al. 2019; Almarri et al. 2020), consistent with the Arab slave trade. However, the Central, Eastern, and Southern African affinities observed here suggest additional contributions linked to 17th–20th century Red Sea and trans-Saharan slave trades, when Eastern (e.g. Ethiopia, Eritrea, Somalia, and Sudan) and Central (e.g. Chad and Congo Basin) African regions were major sources of enslaved individuals (Miran 2022). These historical processes are reflected in elevated effective population sizes (**Figure 4**) and reduced ROH burdens (**Figure 3**) in the most admixed Saudi clusters.

Against this backdrop of intercontinental mobility, persistent consanguineous and endogamous practices have maintained substantial reproductive isolation among clans and tribes, generating genetically distinct sub-clusters in close geographic proximity, consistent with previous reports (Mineta et al. 2021). Although recent admixture with Africans, Europeans, and Central/South Asians may continue, its spread is likely constrained by social structure, as suggested for the neighboring Emirati populations (Elliott et al. 2022). Endogamy and consanguinity increase runs of homozygosity (SROH), elevating health risks. Consistent with high consanguinity rates reported in Madinah and the Western region (El-Mouzan et al. 2007), we observe pronounced signatures of endogamy in Western clusters (clusters 4, 5, 7, and 10), including longer ROHs, reduced effective population sizes, and limited admixture (**Figure 3**). These practices likely counteract the diversity-enhancing effects of admixture, promoting reduced diversity and enrichment of functionally consequential alleles.

Demographic history is known to influence the pattern of variation, particularly for deleterious variation. Classical bottleneck models predict depletion of rare variants and enrichment of low-frequency deleterious alleles due to drift. While this pattern is typical in isolated European populations (Wang et al. 2014; Pedersen et al. 2017; Locke et al. 2019), Saudi genomes display a distinct profile. We detect an excess of rare variants (AF < 1%) relative to gnomAD-AFR, possibly reflecting heterogeneous African-related ancestries (e.g. from Eastern Africa) and/or contributions from Basal Eurasian lineages uncommon outside of Arabia, though we were unable to disentangle between the two due to their collinearity (**Figure S16**). Because both lineages harbor ancestry that predates or bypassed the primary bottleneck of the out-of-Africa expansion, distinguishing them requires either ancient Arabian genomes from the period of Basal Eurasian isolation, which do not yet exist, or reference panels that better resolve Eastern versus Western African ancestry components; the latter is more tractable. Beyond the excess of rare variants, deleterious alleles are nonetheless enriched within the 0.5–5% frequency range. This pattern likely reflects the combined effects of historical bottlenecks and persistent endogamy. Although increased homozygosity may theoretically facilitate purging, reduced effective population sizes limit the efficiency of purifying selection. Subgroups with higher endogamy show greater enrichment of deleterious alleles than more admixed clusters (**Figure 5B**), consistent with findings from other bottlenecked populations (Lohmueller et al. 2008; Lim et al. 2014; Locke et al. 2019) and supporting prior evidence of limited genetic purging in Saudi Arabia (Greater Middle East Variome Consortium et al. 2016).

We emphasize that sub-clusters were defined based on genetic similarity and used as analytical units due to limited tribal affiliation data. In admixed populations, such clustering may obscure finer-scale structure, particularly when distinct Arabian-origin groups share external admixture sources. Although our clusters broadly correlate with geography and show partial concordance with previously described indigenous tribal groupings (Mineta et al. 2021), they should be interpreted as genetically similar groups rather than direct representations of tribal lineages. Now with a better understanding of the genetic history of Saudi Arabians, future studies incorporating improved demographic metadata and ancestral reference panels may refine these inferences using ancestry-specific approaches (Moreno-Estrada et al. 2013; Browning et al. 2016; Tang and Chiang 2025).

These limitations notwithstanding, collectively, our results demonstrated that Saudi Arabia’s demographic history has shaped its contemporary genetic landscape with potential implications for health. The persistence of endogamy and consanguinity continues to elevate genetic risk and remains common (Warsy et al. 2014; Albanghali 2023). Public health initiatives, including mandatory premarital screening and genetic counseling (Saffi and Howard 2015; Delatycki et al. 2020; Aleissa et al. 2022), have increased awareness and informed reproductive decision-making. Although such programs have not substantially reduced at-risk marriages, they have contributed to reductions in affected births through prenatal diagnosis and related interventions (Saffi and Howard 2015; Albanghali 2023; Khayat et al. 2024). Continued education and socioeconomic development may further influence the prevalence of consanguineous unions, which remain more common in rural and less-educated communities (Tadmouri et al. 2009).

Finally, our study helps address the persistent under-representation of Arabian populations in global genomic datasets (Greater Middle East Variome Consortium et al. 2016; Almarri et al. 2020; Elfatih et al. 2024; Oleksyk et al. 2025). In gnomAD, individuals of Middle Eastern origin comprise only 0.2% of genomes and 0.38% of exomes, compared to substantially higher representation of European and African populations (e.g. 44.6% and 27.3% of genomes). This imbalance limits clinical knowledge and application of regionally enriched variants (Greater Middle East Variome Consortium et al. 2016) and reduces the accuracy of polygenic prediction and imputation in Arabian populations (Thareja et al. 2021; Cahoon et al. 2024). Notably, 9.7% of protein-altering variants identified here are previously unreported and disproportionately predicted to be deleterious. Expanding representation of Middle Eastern populations in reference panels such as gnomAD and TOPMed is therefore essential (Oleksyk et al. 2025). Acknowledging that public data sharing requires careful consideration of national and community interests, we provide allele frequencies for 25,488,981 high-quality variants to enhance genomic resources for this underrepresented population.

## METHODS

### Data collection, processing and quality control

For all studied samples, written informed consent was obtained from each participant, all of whom were above 18 years of age. The human subjects for this study were derived from a comprehensive collection of Institutional Review Board (IRB)-approved research protocols focused on the genetics of various diseases and control groups. These protocols, approved under the Saudi Genome Project Satellite site, include approval numbers 16-300, 16-310, and 20-211, and were reviewed and approved by the IRBs at King Abdulaziz City for Science and Technology (KACST) and King Fahad Medical City (KFMC). Tribal affiliations reported by some participants were anonymized and referenced solely by their geographical locations. In compliance with Saudi privacy legislation and the protection of human subject confidentiality, the sharing of raw genotyping and clinical data is restricted. Access to this data requires prior approval from the Saudi National Bioethics Committee.

#### Array data

##### Sample collection, genotyping and quality control

A total of 3,752 samples were collected in Saudi Arabia between the years 2017 - 2020 as control individuals for various projects, such as the GenOMICC International project and covid19 host genetics consortium studies (COVID-19 Host Genetics Initiative et al. 2022; Pairo-Castineira et al. 2023). Individuals were genotyped on the Axiom Genome-wide CEU 1 Array including customized variants following the manufacturer’s specifications for sample preparation, including whole genome amplification, fragmentation, denaturation, and hybridization. Genome-wide SNP genotyping was performed using the automated, high-throughput GeneTitan system from Affymetrix.

We filtered individuals with sample call rates < 0.9 using PLINK v1.9 (Chang et al. 2015) on each plate individually before merging the autosomal SNPs across the different plates, resulting in a merged set of 757,790 SNPs. We removed duplicates and non-biallelic variants, retaining 703,986 SNPs. We then filtered SNPs with greater than 10% missing rate and SNPs that did not pass Hardy Weinberg Equilibrium (HWE) test (*P* < 10^-6^) using PLINK, resulting in a total of 606,349 SNPs for analysis. We lifted over the genomic coordinates from human reference genome hg19 to hg38. We phased the data using Beagle v5.2 (Browning et al. 2021).

##### Removing close relatives and filtering out outliers

Using the 3,752 Saudi samples and 606,349 SNPs, we pruned the dataset by linkage disequilibrium (LD) (using the command *--indep—pairwise 50 5 0.8* in PLINK), resulting in 547,307 SNPs to estimate individuals’ relatedness using King v2.2.5 (Manichaikul et al. 2010). We removed twins (or duplicated individuals) as well as first degree relatives, retaining 3,403 samples. Furthermore, we performed PCA and performed two iterations of outlier (defined as being > 6 standard deviation (SD) away from the mean in any of the first 10 PCs), resulting in 3,352 samples left for further analyses.

##### Defining and imputing the samples’ regional affiliation

Due to privacy protection and ethical restrictions, we did not have access to specific tribal name of each individual to help define the unit of analysis in our study or help interpret our findings. Instead, we have limited information of the geographic region of origin of the participant, inferred based on their family name that is linked to geographical location. We thus aimed to use available regional geographical affiliations to validate and interpret the results of clustering based on genetic data. However, 82% of the individuals in our data (2,740 of the 3,352) did not have self-reported geographical information. We thus imputed such information using the software HARE (harmonized ancestry and race/ethnicity) package (Fang et al. 2019) based on the available Self-identified Race/Ethnicity (SIRE) regional information of 612 individuals. SIRE in our data were derived from either self-report or individual’s family name that is presumed to reflect their tribal affiliation linked to a particular geographical location (**Table S1**). The HARE package combines genetically inferred structure based on PCA with available SIRE information to train a support vector machine (SVM) classifier that could correct for potentially mislabeled SIRE and predict the race/ethnicity, in this case geographical regional label, for those individuals missing SIRE. We used the HARE to impute regional information of the samples missing a SIRE label in our dataset using the first 30 PCs as the input data. We used the highest predicted membership probability (L_1_) labels to aid in the interpretation of the population sub-clusters that we infer from genetic data.

#### Whole genome sequencing (WGS) data

##### Sequencing information and processingh4

In addition to the genotyped samples, 349 samples were whole-genome sequenced (WGS) to a targeted depth of 30x. The samples were prepared following the Illumina’s TruSeq Nano sample preparation protocol and sequenced on an Illumina HiSeq X-ten machine. The raw sequences were aligned against the human reference genome GRCh38 using the Burrows-Wheeler Aligner (BWA) version 0.7.10 (Li and Durbin 2009). Picard tools version 1.117 was used to mark duplicates (McKenna et al. 2010). All sample preparation, sequencing, sequence alignment, pre-processing, quality control before calling of variants and BAM file augmentation were performed by deCODE genetics (https://www.decode.com), and a more detailed information on these steps is documented in Jónsson et al. (Jónsson et al. 2017).

##### Variant calling and filtering

We merged the gVCFs of the 349 samples using CombineGVCFs in GATK (Van Der Auwera et al. 2013) and subsequently performed a joint genotyping calling using GenotypeGVCFs. We performed variant quality score calibration (VQRS) on the combined samples using VariantRecalibrator and ApplyVQRS in GATK (Van Der Auwera et al. 2013). We supplied the homo sapiens reference assembly 38 (Homo_sapiens_assembly38.fasta) and used the following resources: HapMap III variants were used as training and truth sets with prior priority of 15, 1000G omni2.5 sites were used as training set with prior priority of 12, 1000G phase1 high confidence SNPs was used as training set with prior priority of 10 and the dbSNP138 as known SNPs with prior probability of 2. For the annotations, we included the QD, MQ, MQRankSum, ReadPosRankSum FS and SOR. We used 99% sensitivity level to filter the SNPs.

##### Quality control on samples and SNPs

All 349 samples had missing genotyping rate < 10%. We excluded 302,640 SNPs with missing rate > 10% and 53,981 SNPs based on HWE threshold (*P* < 10 ^-6^), leaving 26,781,476 SNPs. We removed non-biallelic sites which left 26,408,559 variants. Further filtering was applied on specific downstream analyses when appropriate. To exclude outliers in our samples, we merged the 349 WGS samples with our array data and the HGDP dataset at segregating SNPs shared across all datasets. A principal component analysis (PCA) was performed using PLINK and we used HARE to impute missing self-reported individual nationalities (e.g. self-identified nationality as Saudi or not). We excluded 8 samples which were not imputed as a Saudi. We then removed monomorphic sites which were introduced by calling the variants including these potentially non-Saudi samples, leaving 25,488,981 variants.

We filtered samples based on relatedness using King software v2.2.5 (Manichaikul et al. 2010). For estimating the relatedness, we randomly sampled 550,000 SNPs with minor allele frequency > 1% after LD pruning (*--indep-pairwise 50 5 0.5* using PLINK) to estimate the relatedness. We removed 37 twins/duplicates and first-degree relatives. Using the PCA, we further removed 2 samples that appeared as extreme outliers (> 6 SD on any of the first 10 PCs), leaving 302 samples. Haplotype phasing was performed on the remaining 302 samples and 25,488,981 variants using Eagle v2.4.1 (Loh et al. 2016).

#### Annotation of variants

We annotated the variants using the popular VEP (v.110) (McLaren et al. 2016) as well as two recently published annotation tools, AlphaMissense (Cheng et al. 2023) and Genomic Pre-trained Network (GPN) (Benegas et al. 2023). The AlphaMissense only annotates missense variants and has three functional classes, “likely pathogenic”, “ambiguous” and “likely benign”. The GPN annotates all genomic variants and assign a deleteriousness score to each variant in gnomAD (v3). We downloaded the pre-computed scores from https://huggingface.co/datasets/songlab/gnomad/resolve/main/test.parquet, accessed 2/9/2024.

#### Merging of Saudi whole-genome-sequence data with ancient genomes

We first back-converted the genomic coordinates of the Saudi whole-genome sequencing (WGS) data from the hg38 reference assembly to hg19 using LiftOver. We filtered the dataset using PLINK v1.9-beta7 (--snps-only, --geno 0, --allow-no-sex) to retain only biallelic SNPs with a 100% genotyping rate. This filtering step removed 725,421 sites containing missing data from the baseline dataset. We then converted the filtered dataset into EIGENSTRAT format and merged the Saudi WGS data with the Allen Ancient DNA Resource (AADR) v.62.0 Eigenstrat files with the Eigensoft mergeit function which merges two data sets into a third producing the union of the individuals and the intersection of the SNPs in the first two. We then merged this dataset with the Eigenstrat genotype data from Soqotra resulting in a final dataset of 1,032,250 SNPs and 18,017 individuals. We generated an additional dataset merging the Saudi WGS and Soqotra Eigenstrat genotype data with the AADR v.62.0 modern human origins dataset, retaining 506,776 SNPs and 22,333 individuals. A complete list of the individuals and their associated group labels for each of the downstream analyses can be found in **Table S14**.

#### Population genetic analyses

We used the larger collection of Saudi genotyped samples to investigate the genetic substructure and historic admixtures of the population. We then utilized the high-density genome-wide marker information from the WGS data to investigate differences in genetic ancestries with aDNA, population size trajectories, and allelic architecture of functional variants within the social structure of the Saudi population.

##### Evaluation of population structure and identifying discrete population clusters

We merged the fully filtered array and WGS datasets, based on segregating markers. We performed PCA followed by UMAP (McInnes et al. 2018; Diaz-Papkovich et al. 2019) to combine the first 10 PCs and reduced them into two-dimensions in order to explore the population structure. Based on the UMAP results, we assigned individuals to subpopulations using K-means clustering from the R package *stats*. To determine the optimal number of K clusters, we used the Average Silhouette Width (ASW), which is a popular and trusted method to produce quality clustering (Rousseeuw 1987; Batool and Hennig 2021). The ASW uses values between -1 and 1 to measure how similar/dissimilar is an object to others within its cluster as well as objects in different clusters, with higher numbers representing a better fit and appropriateness of clustering. Likewise, a high ASW value corresponds to an optimal number of K clusters for partitioning a particular set of objects. Based on ASW, we determined that K = 12 is the optimal value that best fit the data, although K = 5 or 9 could be equally sensible (**Figure S18**). We validated these clustering by evaluating the concurrence between the clusters and the tribal region assignments. We used these clusters as representative of the social structure and also used them in the whole genome sequencing samples to evaluate patterns of genetic diversity within the Saudi population.

##### Analysis of ancestry components

We conducted the unsupervised admixture analysis using ADMIXTURE software v1.3 (Alexander et al. 2009). We conducted 10 independent runs of admixture analysis for each K and retained the run with maximum likelihood. We used the cross-validation procedure, implemented in the program, to identify the best number of ancestral populations K which fits our data (**Figure S3C**).

##### Saudi and ancient genome analyses

We focused on investigating the genomic history and genetic ancestry of the Saudi, leveraging the available ancient DNA data in AP and surrounding region. We utilized multiple formulations of the D and *f*-statistics (both *f_3_* and *f_4_* statistics) as well as admixture modeling from qpAdm. All of these analyses were performed using the functions in the admixtools R package.

##### D-statistics

We computed two configurations of D-statistics to assess genetic affinity and symmetry of Saudi clusters to four ancient Arabian populations. We filtered results to include only statistics based on at least 50,000 SNPs. For both analyses, the parameters for the qpdstat function include f4mode = FALSE, boot = FALSE and allsnps = TRUE. We performed two specific configurations of the D-statistic:

1. D-symmetry: To test for cladality, we computed statistics of the form D(Test, SaudiCluster; WorldPop, Karitiana.DG), where “Test” represented either ancient Bahrain or Soqotra, “WorldPop” comprised diverse ancient and present-day Eurasian and African populations, and for the outgroup population we selected Karitiana, an indigenous Brazilian population following Sirak et al. (Sirak et al. 2024). Significantly negative D-statistic (Z-score < -3) estimates indicate greater shared drift between WorldPop and Saudi clusters relative to the ancient Arabian Test populations, while significantly positive values demonstrate the inverse relationship, with either result rejecting the hypothesis of population continuity.
2. D-affinity: To identify populations sharing excess genetic drift with Saudi clusters relative to ancient Arabia, we computed statistics of the form D(WorldPop, Test; SaudiCluster, Karitiana.DG). Significantly negative D-statistics indicate that Saudi clusters share more excess drift with the Arabian Test population than with WorldPops. Conversely, significantly positive values denote the inverse: greater shared drift between Saudi clusters and WorldPops.

##### *f*4-statisitcs for Basal Eurasian ancestries

We used two approaches to estimate Basal Eurasian ancestry. First, we used the *f*4-statistic of form *f*4(TestPops, Han.DG, Russia_UstIshim_IUP_snpAD.DG, Outgroup) and implemented this via the qpdstat function with f4mode = TRUE. The TestPops included the Saudi clusters and a selection of present-day Middle Eastern, Arabian, and East African populations, while Yoruba.DG, Saharawi.DG, Mozabite.DG, and Morocco_Iberomaurusian.AG served as the outgroups (**Figure S8A, S8B**; **Table S12**). Second, on an individual level within the Saudi clusters we modeled the relationship between African and Basal Eurasian ancestry and genomic diversity using a linear mixed-effects model (LMM) and a nested model comparison via likelihood ratio test. For these model comparisons, Basal Eurasian ancestry proportions were estimated in each Saudi individual using f4-ratio statistics via the admixtools2 package, with the ratio defined as f4(Saudi, WHG; UstIshim, Kostenki14) / f4(Iberomaurusian, WHG; UstIshim, Kostenki14). These estimates were merged with African ancestry components (K=4 in ADMIXTURE analysis) and observed heterozygosity (O{HET}) calculated from phased whole-genome sequencing (WGS) data via vcftools. Prior to modeling, both continuous ancestry predictors (f4-ratio and K4) were z-score standardized (mu = 0, sigma = 1) to allow for direct comparison of their effect sizes (standardized beta coefficients).

#### Linear Mixed-Effects Modeling

To model the relative impact of Basal and African ancestries while accounting for Saudi substructure, we fitted a Linear Mixed-Effects Model (LMM) using the lme4 and lmerTest packages. Ancestry components were treated as fixed effects, and Saudi cluster membership was included as a random intercept. P-values for the fixed effects were estimated using Satterthwaite’s degrees of freedom method. To assess the impact of correlation between the two ancestries we calculated the Variance Inflation Factor (VIF) using the car package in R. We evaluated model performance using Nakagawa and Schielzeth’s approach to partition variance into Marginal R^2^ and Conditional R^2^ where the former estimates the proportion of total variance explained exclusively by the fixed ancestry effects, while the latter measures the variance explained by the entire model (fixed ancestries + random clusters). To evaluate the unique contribution of each ancestry, we calculated the Partial R^2^ via nested model comparison. This was achieved by subtracting the Marginal R^2^ of a reduced model (lacking the predictor of interest) from the Marginal R^2^ of the full model. This metric isolates the percentage of heterozygosity variance strictly attributable to a specific ancestry, independent of the other. Finally, we adjusted for between cluster variance by computing within-cluster partial correlations using the ppcor package. Here we residualized the observed heterozygosity, f4-ratio Basal Eurasian ancestry estimates and ADMIXTURE African ancestry estimate against only the Saudi cluster random intercept and calculated the Pearson correlation between these residuals.

To formally test whether the addition of Basal Eurasian or African ancestry significantly improved the model’s fit, we performed Likelihood Ratio Tests (LRT) using an Analysis of Variance (anova) in R. The significance of model improvement was derived from a chisq distribution of the difference in deviance (2 times log-likelihood) between the Base Model (one ancestry) and the Full Model (two ancestries). We additionally evaluated the Akaike Information Criterion (AIC) to compare relative model quality.

##### *f*_3_-statistics

We computed two forms of *f*_3_-statistics, the outgroup *f*_3_ and admixture *f*_3_, using the Admixtools2 qp3pop function to further investigate genetic affinity between Saudi cluster and ancient samples. To quantify the shared genetic drift between Saudi clusters and diverse set of 1698 Eurasian and African populations dating between the present-day and 50 thousand years ago, we computed the outgroup *f*_3_-statistic of form *f*_3_(Han.DG; SaudiCluster, WorldPop), where the Han.DG population served as the outgroup to anchor the comparison. Higher *f*_3_-statistic values in this configuration indicate increased shared genetic drift between the Saudi cluster and the test population relative to the outgroup. We merged the statistical results with metadata to associate each test population with its mean sample age (BP) and geographic coordinates. We generated geospatial heatmaps using ggplot2, plotting test populations on a world map background. Point colors representing *f*_3_-statistic magnitude were mapped using the “Zissou1” continuous palette from the wesanderson package. To analyze temporal trends, the data were stratified into four time periods (T1: <1,000, T2: 1,000–3,000, T3: 3,000–10,000, and Distal: >10,000 BP) and faceted by Saudi cluster.

To test whether Saudi clusters are the result of admixture between specific ancient Arabian populations and diverse global groups, we computed the admixture *f*_3_ statistic of form *f*_3_(SaudiCluster; Test, WorldPop), where “Test” was either ancient Bahrain or Soqotra, and “WorldPop” included diverse Eurasian and African populations. Negative *f*_3_ statistic values with significant Z-scores (Z < -3) were interpreted as evidence that the Saudi cluster is a mixture of lineages related to the Ancient Arabian Source and the WorldPop Population. We visualized significant admixture signals using “flight-path” maps generated with ggplot2, sf, and rnaturalearth. For the subset of four target Saudi clusters that produced significant *f*_3_ statistic values (c3, c8, c11, and c12), we plotted curved lines connecting the fixed Ancient Arabian Source to the Global Population.

##### Ancestry Modeling with qpAdm

We performed temporal qpAdm admixture modeling to reconstruct the ancestry of Saudi clusters through time and space. To form a set of consistent candidate source populations for qpAdm analyses we utilized a series of initial outgroup *f*_3_ and PCA analyses. Specifically, we repeated our outgroup *f*_3_-statistic test as described above with a pooled Saudi population consisting of individuals from each of the 12 clusters. From the *f*_3_-statistic test results we stratified the Test populations into three time periods (T1 = <1000, T2 = [1000 - 3000], and T3 = >3000 BP) and the following 13 global regions: Eastern Europe, Western Europe, Southern Europe, Caucasus, Levant, Iran, Anatolia, Arabia, Central Africa, Western Africa, Northern Africa, Southern Africa, and Eastern Africa. Within each time period, we merged into meta-population source for downstream qpAdm analysis those populations that had similar outgroup *f*_3_-statistic estimates at the top of the range and also displayed evidence of clustering in principal component analysis (PCA) space (**Figure S19, Table S14**). PCA analysis was performed with the smartpca function in EIGENSOFT v.7.2.0 using the parameter file options, numoutlieriter: 0, numoutlierevec: 0, numoutevec: 10, newshrink: YES, hiprecision: YES, and lsqproject: YES, and projecting the ancient genomes genotyped on the Affymetrix Axiom Genome-Wide Human Origins 1 array (“HO” AADR v.62.0) onto 70 global modern populations and the Saudi cluster individuals. The list of meta-population sources derived from AADR can be found in **Table S14**, with their age distributions visualized in **Figure S19**.

We then performed qpAdm across four distinct time periods: the three proximal time-stratified models (T3, T2, and T1) plus a distal model (regional samples older than T3). For all models, we utilized the parameters allsnps = TRUE and fudge_twice = TRUE.

##### Distal Modeling

For the Distal model, we employed a “rotation” strategy where no populations were fixed in the reference (right) set. Instead, we rotated a pool of nine source populations between the target (left) and reference (right) positions. The rotating sources were: CHG, Morocco_Iberomaurusian.AG, Greece_Crete_Aposelemis_N.AG, Turkey_Central_Boncuklu_PPN.AG, Turkey_Southeast_Cayonu_PPN.SG, Iran_GanjDareh_N.AG, Israel_Natufian.AG, WHG, and EEHG. We tested all possible combinations of 1 to 5 sources.

##### Proximal Modeling (T3, T2, T1)

For the time-stratified models, we used a fixed set of outgroup (right) populations: CHG, Morocco_Iberomaurusian.AG, Greece_Crete_Aposelemis_N.AG, Iran_GanjDareh_N.AG, Turkey_Southeast_Cayonu_PPN.SG, WHG, EEHG, Israel_Natufian.AG, and Turkey_Central_Boncuklu_PPN.AG. We performed a combinatorial testing of 1 to 6 sources from specific candidate pools for each period. Due to the challenge of controlling for qpAdm model violations resulting from left and right-set admixture (Flegontova et al. 2025), especially with modern target and ancient source populations (Agranat-Tamir et al. 2020), we restricted our proximal modeling to a nonrotating protocol using the following sources:

- <u>T3 Candidate Sources:</u> T3_Caucasus, T3_Iran, T3_wEurope, T3_Levant, T3_Anatolia, T3_sAfrica, T3_eAfrica, T3_nAfrica, T3_eEurope, T3_cAfrica, and T3_sEurope.
- <u>T2 Candidate Sources:</u> T2_nAfrica, T2_Anatolia, T2_Iran, T2_eAfrica, T2_sAfrica, T2_Caucasus, T2_wEurope, T2_Levant, T2_eEurope, T2_sEurope, and three ancient Arabian sources (T2_Arabia_EMTylos_SeleucidCharacene.SG, T2_Arabia_LTylos_Sasanian_MH1MH2.SG, T2_Arabia_LTylos_Sasanian_MH3.SG).
- <u>T1 Candidate Sources:</u> T1_Iran, T1_eAfrica, T1_Caucasus, T1_sEurope, T1_sAfrica, T1_Levant, T1_eEurope, T1_cAfrica, T1_wEurope, T1_Anatolia, T1_nAfrica, and T1_Arabia.

We filtered results to retain only plausible models, defined as those with valid admixture weights [0-1] and p-value ≥ 0.01. For each target population, we selected the two plausible representative models based on the highest p-values and least number of sources (**Figure 2C-D**). Error bars representing standard errors were included for all admixture components. In addition to the qpAdm model *p*-value, we also reported the *p*-value testing whether the difference between two models of rank difference 1 is significant (nested *p*-value) in **Table S8-S11**. Only the T2 and the Distal modeling produced plausible models based on these criteria, suggesting that in T1 and T3 we may not have adequate ancestry references for modeling.

##### Runs of homozygosity

ROH are continuous segments of homozygous genotypes inherited from common ancestor (Ceballos et al. 2018). Following Choudhury et al. (Choudhury et al. 2020), we used PLINK function --*option-homozyg* to identify runs of homozygosity (ROH) using the following parameters: we considered at least 100 SNPs for ROH, with a total length ≥ 100 kilobases and at least one SNP per 50 kb on average; we set a scanning window to contain 100 SNPs, allowed 1 heterozygous call and 5 missing calls per scanning window. We used three component Gaussian mixture model from the Mclust package (v.6.1) in R following Pemberton et al. (Pemberton et al. 2012) to classify the ROHs into short (< 635 kb), intermediate (between 635kb and 1671kb) and long (> 1671 kb) sizes. Short ROHs indicate homozygosity from ancient or distant ancestry, i.e. background relatedness. Intermediate ROHs likely arise from background relatedness with moderate level of inbreeding from past few generations, often due to reduced population sizes or reproductive isolation (e.g. due to geographic or cultural preferences), or from recent bottlenecks followed by recovery. Long ROHs arise through recent inbreeding and are common in populations with high levels of consanguinity (Thompson 2013; Ceballos et al. 2018).

##### Demographic history

Utilizing the phased WGS data, we estimated effective population sizes at different time points within the Saudi sub-clusters using RELATE v1.1.9 (Speidel et al. 2019). We used the RelateFileFormats in the Relate package to convert files from VCF format into haps/sample file format. For ancestral allele flipping, we provided RELATE with the human ancestor sequences release 107. We computed the genealogical trees using the parameters -m 1.25e-8 -N 30,000 and subsequently used the EstimatePopulationSize.sh script provided with the Relate package to estimate the effective population sizes. To lessen the bias in inference due to excessive amount of ROH, we performed the analysis by randomly selecting one phased haplotype per individual and ran Relate in haploid mode.

#### Enrichment of functionally deleterious alleles

We compared the allelic architecture between Saudi Arabian and the gnomAD v4 African/ African American (gnomAD-AFR), non-Finnish European (gnomAD-EUR) and Middle Eastern (gnomAD-MID) populations. To check for potential enrichment or purging of deleterious alleles in the Saudi, we computed the ratio of the proportional site frequency spectra for the deleterious alleles in Saudi to gnomAD-AFR or gnomAD-EUR, and contrasted it to the same ratio based on neutral or benign alleles. Utilizing the gnomAD exomes, which have a larger number of Middle Easterners compared to the genomes, we also made comparisons between the Middle Easterns and gnomAD-AFR and gnomAD-EUR. Significance differences in the ratios between variants functional classes were tested through bootstrapping.

For every comparison between populations or subpopulations, we used Hypergeometric (v 3.6.2) distribution in R to down-sample both populations to equal sample sizes. All exome comparisons were down-sampled to gnomAD-MID sample size. To account for technical differences in data generation of WGS call sets between gnomAD and Saudi data, we used the proportions of variants from the normalized allele frequency spectra rather than number of variants to compare the ratio between the Saudi and the gnomAD populations at a given allele count or frequency bin. However, when comparisons were made between two gnomAD populations or between two Saudi subpopulations, the actual number of variants were used.

## DATA AVAILABILITY

In compliance with Saudi privacy legislation and the protection of human subject confidentiality, access to individual and clinical data requires prior approval from the Saudi National Bioethics Committee. The Saudi variants discovered through WGS and their estimated allele frequencies are deposited in the Figshare repository and can be accessed through this link: https://doi.org/10.6084/m9.figshare.28059686.v1 and the array allele frequencies estimated per Saudi sub-cluster can be accessed via https://doi.org/10.6084/m9.figshare.28280060.

## ACKNOWLEDGMENTS

We are very grateful to the study participants who donated the samples used in the study. We would like to thank all the volunteers over the years for their invaluable contributions to this research. Special thanks are extended to Albandari Alowayn, Rasha Aljelaify, Mariam AlSaeed, Amal Almutairi, Fatimah Alqubaishi, and Ebtehal AlSolme for their assistance with sample collection and processing, as well as to Hadeel Elbardisy and Junghyun Jung for their efforts in data processing and preparation. We thank Michael Campbell and Arun Durvasula for providing feedback on the drafted manuscript, and Rui Leite Portela Martiniano for providing us with the Bahrain ancient DNA data.

This study was supported by National Institute of General Medical Sciences (NIGMS) of the National Institute of Health (NIH) under award number R35GM142783 (to C.W.K.C.), the National Institute of Allergy and Infectious Diseases (NIAID) of the NIH under the award number R01AI173172 (to S.M.), the National Science Foundation under the award number 2135954 (to S.M.). This study was also partially funded by King Abdulaziz City for Science and Technology (KACST) (to M.A.), as part of various international genomics health research initiatives, conducted under approved agreements between KACST and the Karolinska Institute, University of Southern California, Brigham and Women’s Hospital, and deCODE Genetics. M.A. has served as the principal investigator for the Saudi Genome Project, funded by KACST, and as the director of a satellite site at King Fahad Medical City between 2016-2023.

We also extend our gratitude to deCODE Genetics and the KACST Genotyping and Sequencing Facilities/Saudi Genome Project for their technical support. Computation for this work was supported by University of Southern California’s Center for Advanced Research Computing (https://www.carc.usc.edu).

## AUTHOR CONTRIBUTIONS

M.A and C. W. K. C. conceived the study. D.K.M., M.P.W. and C.W.K.C. designed the analysis. C. W. K. C., S.M., and M.A. acquired funding for the data generation and analysis in this study. M.A. performed sample acquisition and data generation and processing. D.K.M., M.P.W., L.T., and J.T. analyzed the data. J.T. and C.D.H. provided analysis tools and resources. D.K.M., M.P.W., M.A, C.D.H. and C.W.K.C. interpreted the results. D.K.M., M.P.W. and C.W.K.C. wrote the manuscript with input from all co-authors.

## SUPPLEMENTAL INFORMATION

Word document: Supplementary Figures S1 – S19

Word document: Supplementary Tables S1 – S3, S13

Excel spreadsheet: Supplementary Tables S4-S12, S14

