## Supplementary Figures for "Patterns of population structure and genetic variation within the Saudi Arabian population"

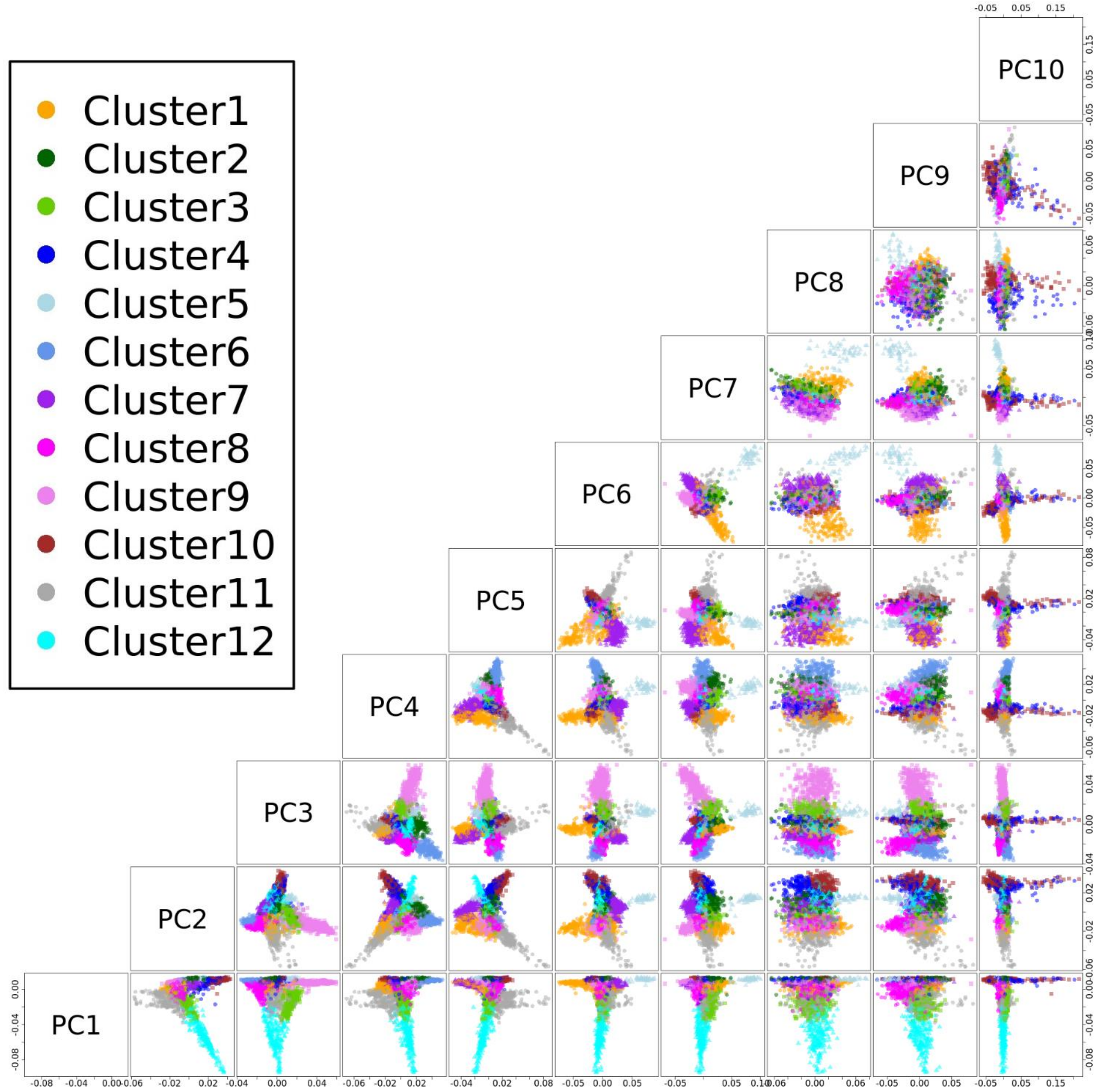


**Figure S1. Principal components analysis of the Saudi Arabian population.** The top 10 PCs are shown in biplots two principal components at a time. Individuals are colored based on the inferred sub-clusters.

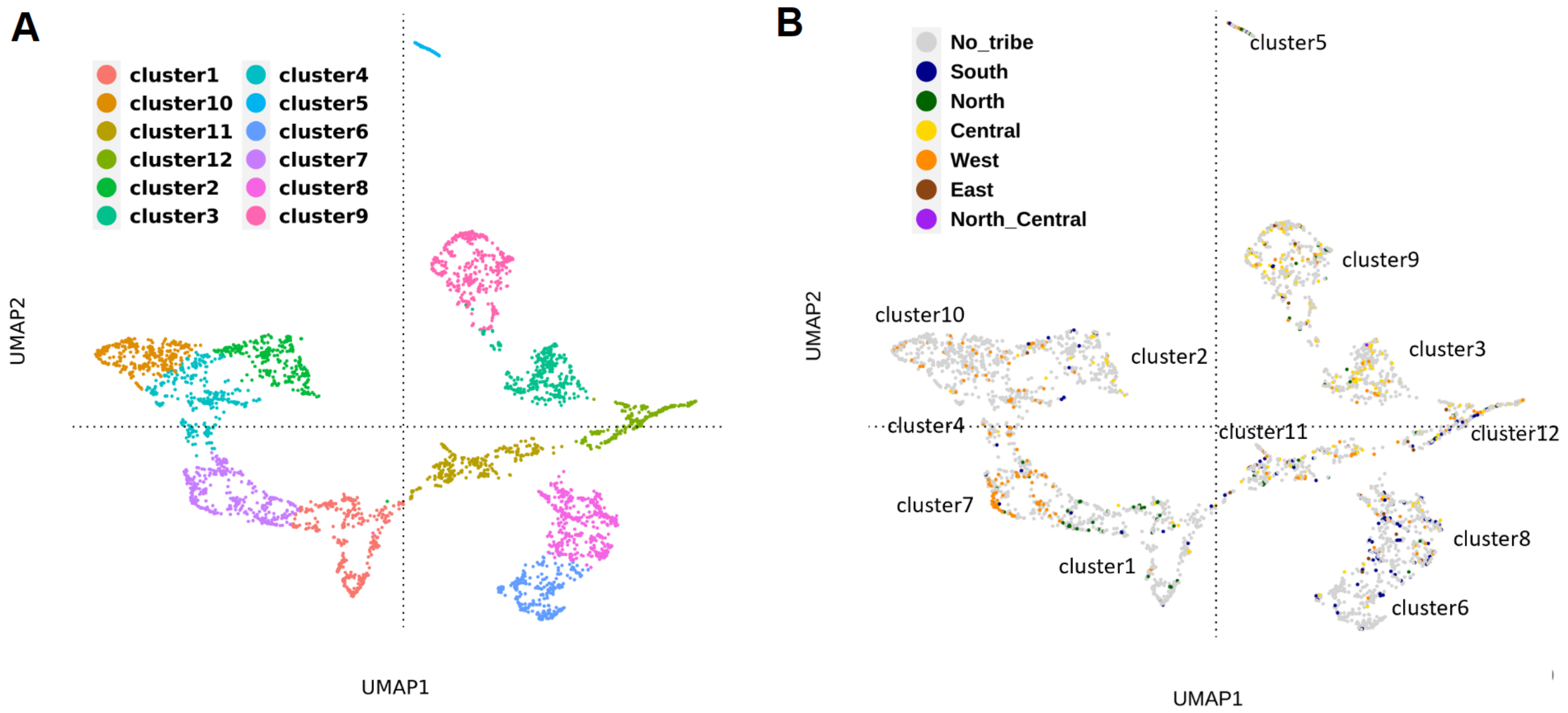


**Figure S2. Average Silhouette Width clustering of Saudi individuals based on the UMAP**. A UMAP of Saudi individuals based on the top ten principal components. In (A), each individual in our dataset was colored by their cluster membership inferred by Average Silhouette Width. In (B), the samples were colored based on the subset of individuals with available self-identified tribal regions. No_tribe refers to the samples without self-identified tribal information. A regional map of Saudi Arabia is included in **Figure 1D** with matching colors to the regional labels in (B).

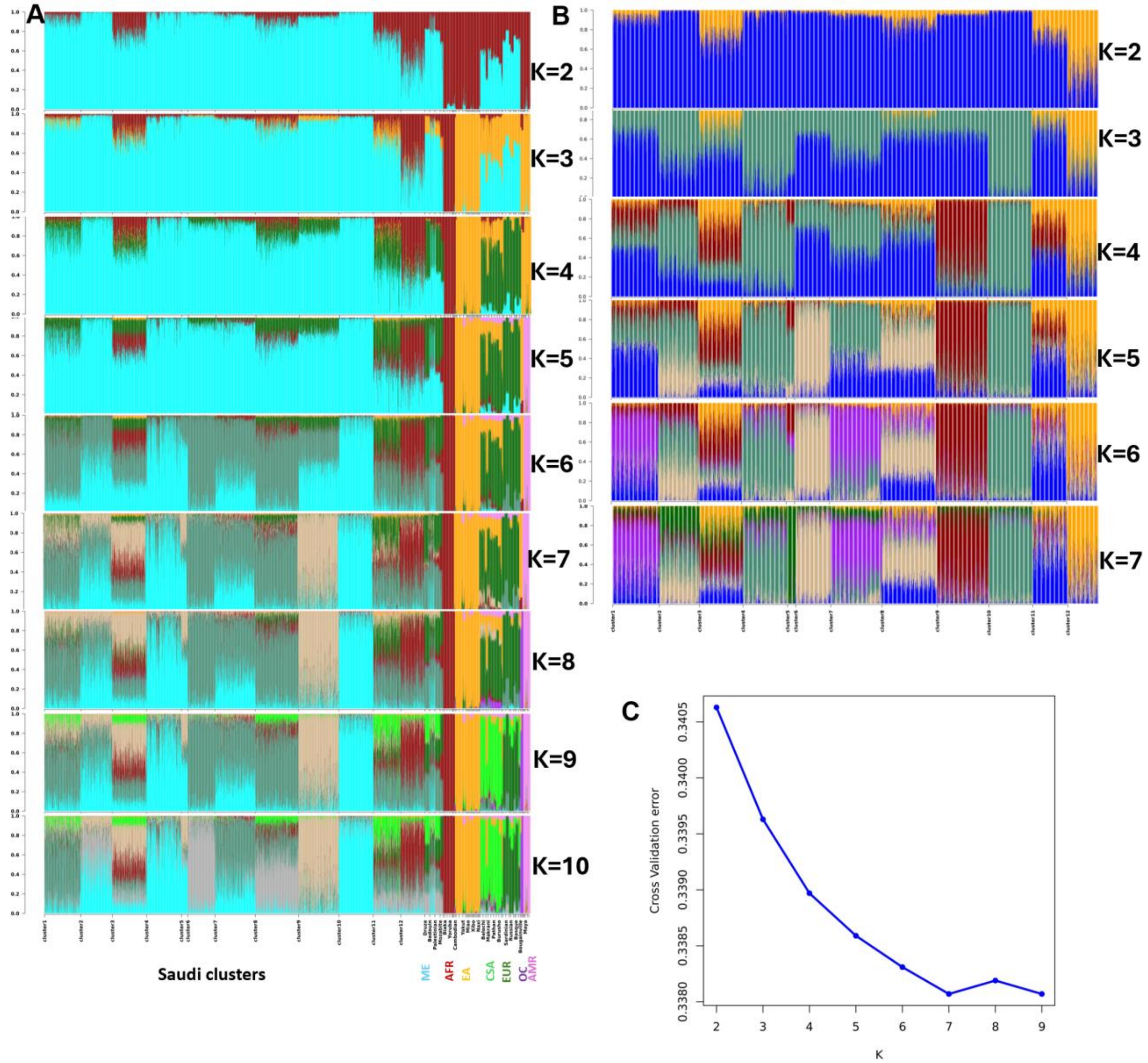


**Figure S3. Admixture analysis.** (A) Admixture analysis for Saudi clusters and HGDP populations for K = 2 - 10. (B) Admixture analysis for Saudi clusters for K = 2 - 7. (C) Admixture cross-validation error estimation for optimal K for Saudi clusters. The K with the lowest cross-validation error was at K = 7, but between K = 5 to 8 new ancestries were introduced only within Saudi samples. New component of ancestries from HGDP reference samples were introduced at K = 9, which we interpreted and discussed in the Main Text. ME – Middle Eastern, AFR – African, EA – East Asian, CSA – Central & South Asian, EUR – European, OC – Oceania, AMR – American. Among the Saudi clusters in (B), cluster5 showed the most homogeneity with a single dominant ancestry component having an average proportion of 94% of the genome, while others e.g. cluster3 showed varying levels of ancestry components with maximum proportion of 27%.

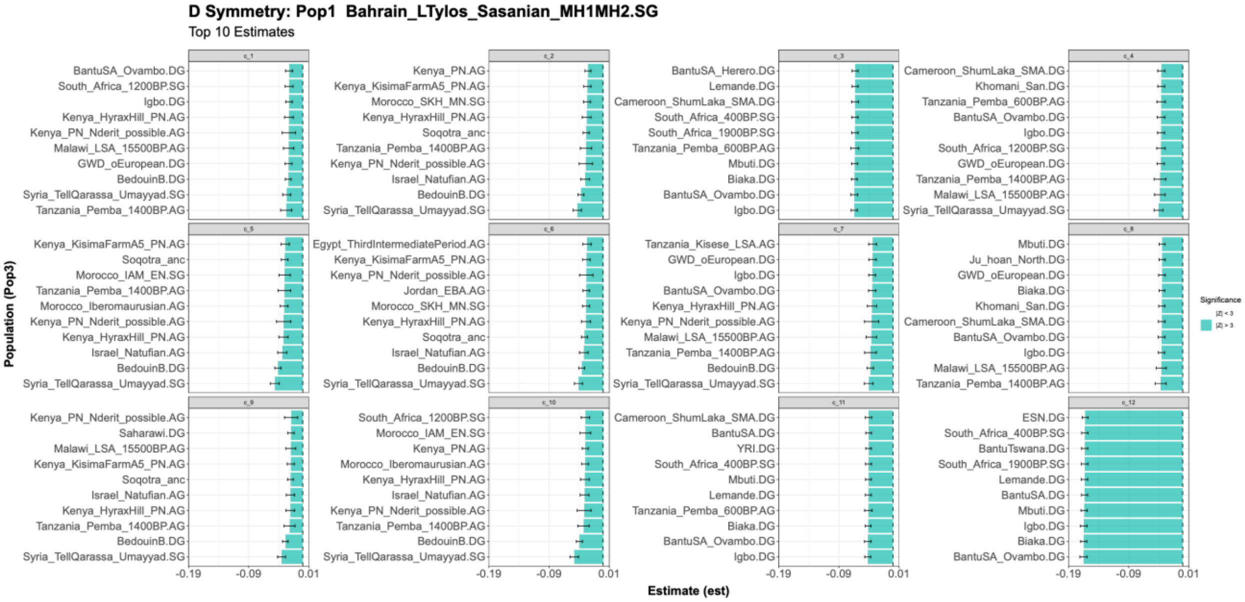
D Symmetry: Pop1 Bahrain_LTylos_Sasanian_MH1MH2.SG
Top 10 Estimates
Population (Pop3)
Estimate (est)
Significance
|Z| > 3

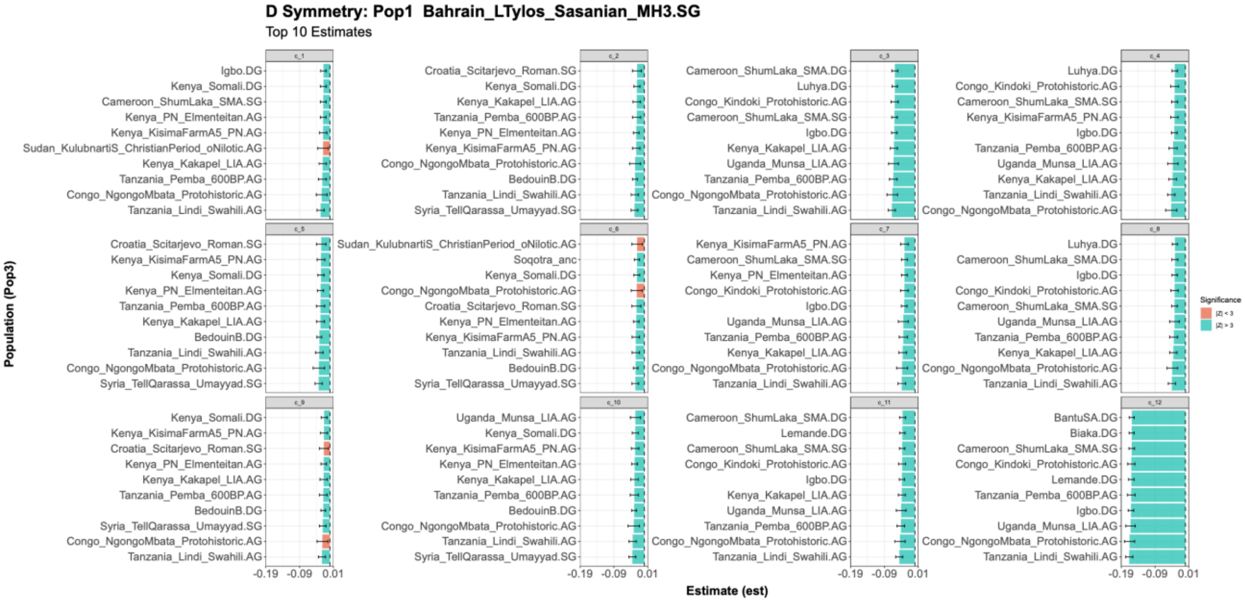
D Symmetry: Pop1 Bahrain_LTylos_Sasanian_MH3.SG
Top 10 Estimates
Population (Pop3)
Estimate (est)
Significance
|Z| < 3
|Z| > 3

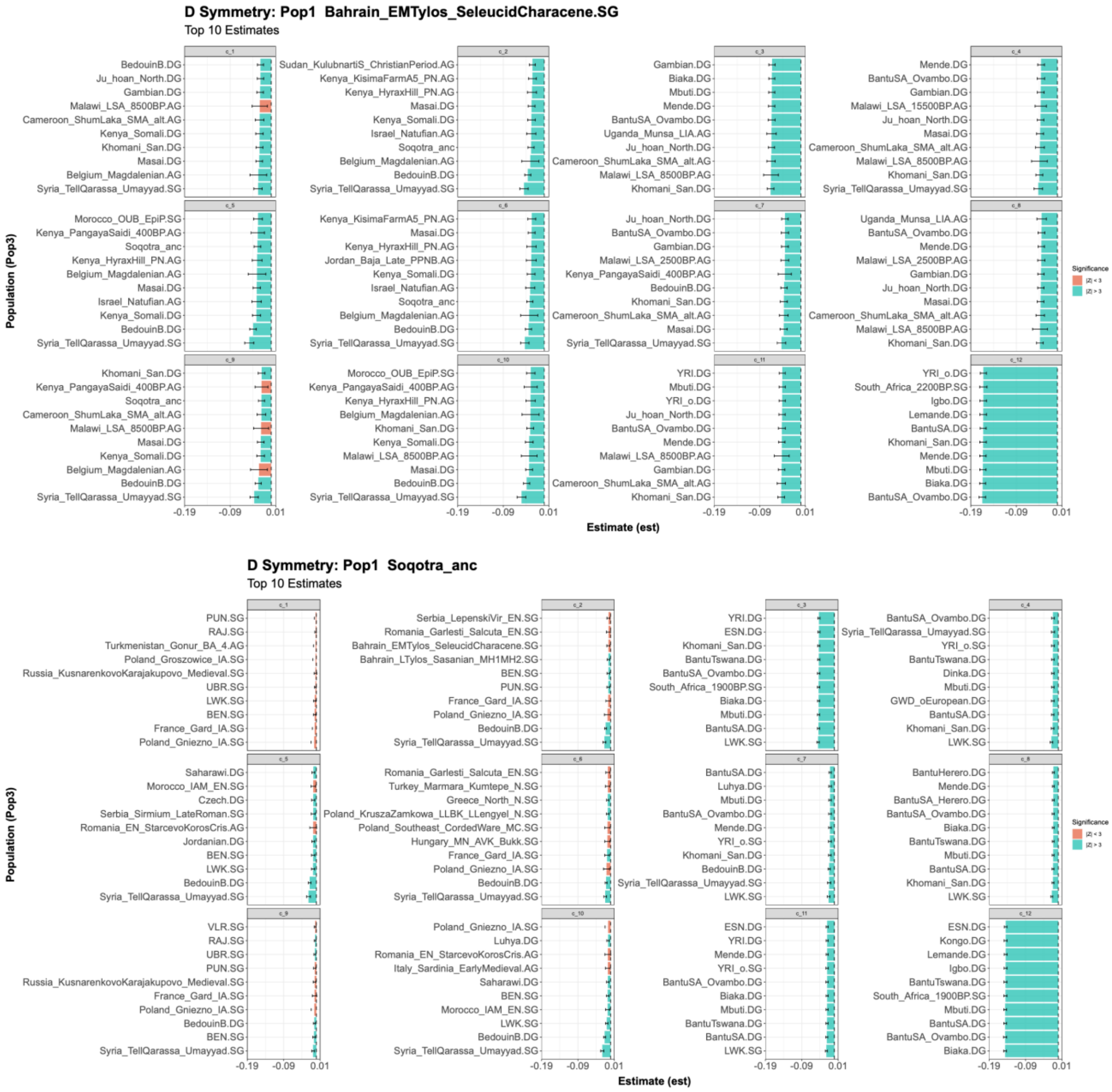


**Figure S4. Symmetry D-statistics to test cladality.** We computed symmetry D-statistics of the form D(Ancient Arabian, Saudi Cluster; WorldPop, Karitiana.DG), where ”Ancient Arabian" represented either ancient Bahrain or Soqotra, "WorldPop" comprised diverse ancient and present-day Eurasian and African populations, and for the outgroup population we selected Karitiana, an indigenous Brazilian population (**Methods**). Significantly negative D-statistic (Z-score < -3) estimates indicate greater shared drift between WorldPop and Saudi populations relative to the ancient Arabian populations, while significantly positive values demonstrate the inverse relationship, with either result rejecting the hypothesis of population continuity. Teal bars indicate statistically significant estimates of shared drift (|Z| > 3), while coral bars denote non-significant results (|Z| < 3). Error bars represent +/- 1 standard error. Across all symmetry

D-statistic tests we reject a model of strict population continuity between ancient Arabian groups and present-day Saudi populations.

We observed a consistent Levantine/Arabian-related deviation via the repeated appearance of Syria_TellQarassa_Umayyad together with BedouinB across all ancient Arabian populations. Other Levantine-related sources also appear to break D-statistic symmetry, namely Israel_Natufian and Jordan_EBA, especially under the Arabian Bahrain MH1MH2 D-statistic test. Further suggestions of heterogeneity amongst the Saudi Clusters emerges from non-African/Near Eastern populations such as Mediterranean/Southern and Eastern-European populations (Croatia_Scitarjevo_Roman/Italy_Sardinia_EarlyMedieval/Poland_Gniezno_IA), especially under the Arabian Soqotra D-statistic test.

Focusing on significantly negative estimates (i.e., increased shared drift with Saudi relative to ancient Arabian tests), we observe a consistent signal dominated by Sub-Saharan African–related populations. Amongst the Sub-Saharan African–related populations many derive from East/Northeast African regions, including Kenya-associated pastoralist/forager/modern proxies (e.g., Kenya_KisimaFarmA5_PN, Kenya_HyraxHill_PN, Kenya_Kakapel_LIA, Kenya_Somali) and coastal/Great Lakes signals (e.g., Tanzania_Lindi_Swahili, Tanzania_Pemba_600BP/1400BP, Uganda_Munsa_LIA).

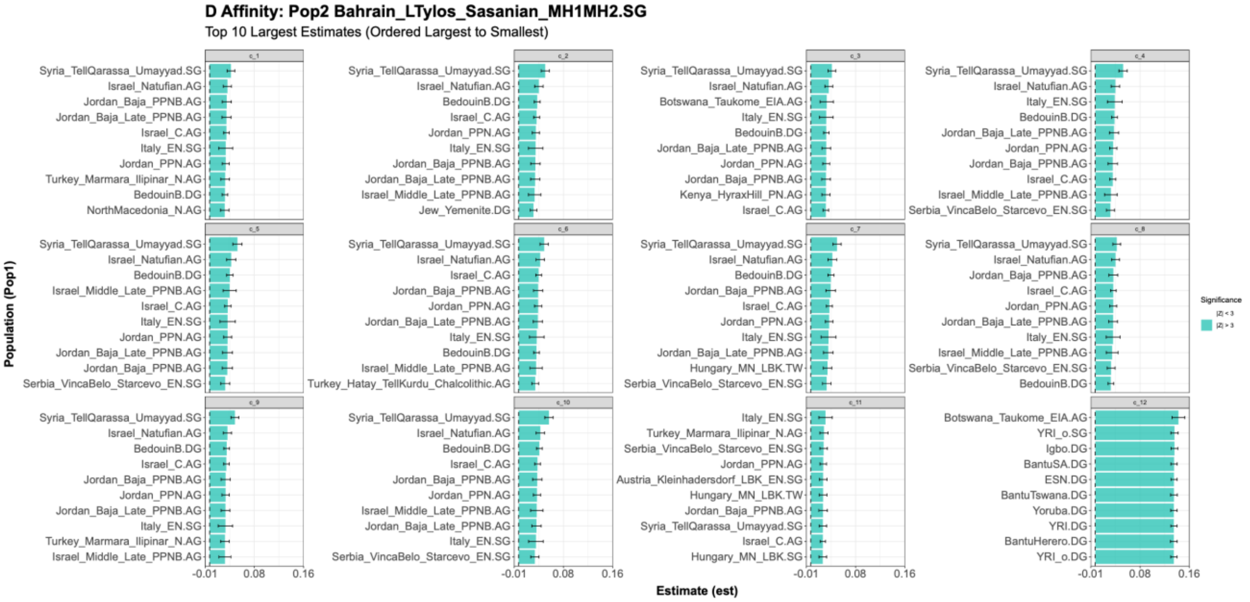
D Affinity: Pop2 Bahrain_LTylos_Sasanian_MH1MH2.SG
Top 10 Largest Estimates (Ordered Largest to Smallest)
Population (Pop1)
Estimate (est)
Significance
|Z| > 3
-0.01
0.08
0.16
Syria_TellQarassa_Umayyad.SG
Israel_Natufian.AG
Jordan_Baja_PPNB.AG
Jordan_Baja_Late_PPNB.AG
Israel_C.AG
Italy_EN.SG
Jordan_PPN.AG
Turkey_Marmara_Ilipinar_N.AG
BedouinB.DG
NorthMacedonia_N.AG
Jew_Yemenite.DG
Israel_Middle_Late_PPNB.AG
Botswana_Taukome_EIA.AG
Kenya_HyraxHill_PN.AG
Serbia_VincaBelo_Starcevo_EN.SG
Turkey_Hatay_TellKurdu_Chalcolithic.AG
Hungary_MN_LBK.TW
Austria_Kleinhadersdorf_LBK_EN.SG
Hungary_MN_LBK.SG
YRI_o.SG
Igbo.DG
BantuSA.DG
ESN.DG
BantuTswana.DG
Yoruba.DG
YRI.DG
BantuHerero.DG
YRI_o.DG

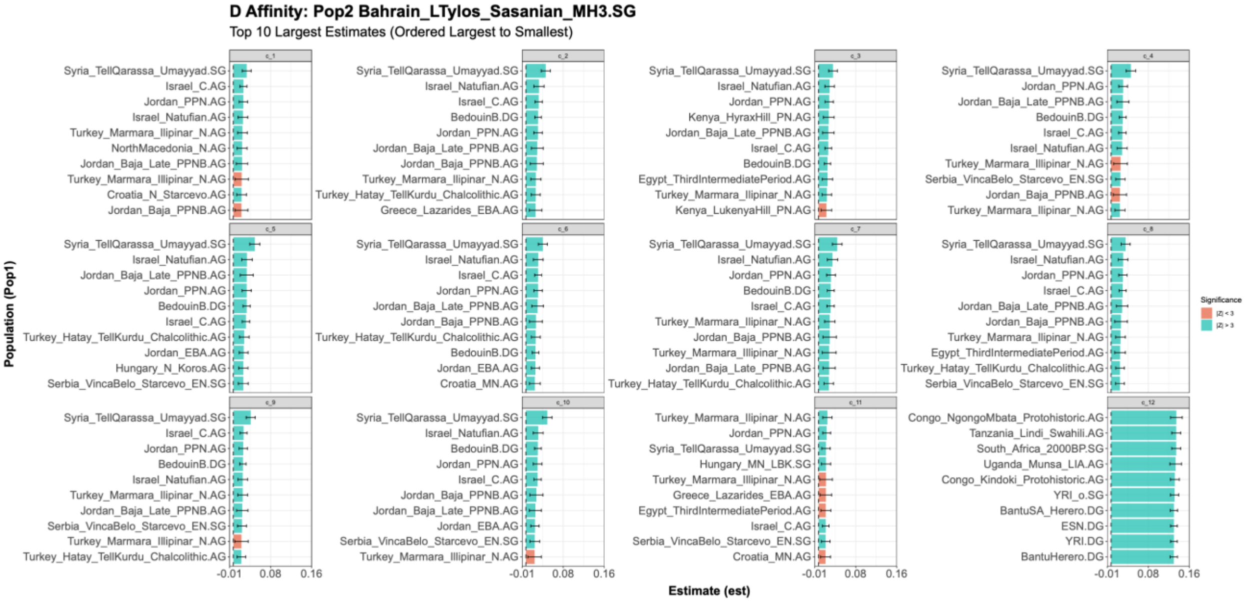
D Affinity: Pop2 Bahrain_LTylos_Sasanian_MH3.SG
Top 10 Largest Estimates (Ordered Largest to Smallest)
Population (Pop1)
Estimate (est)
Significance
|Z| < 3
|Z| > 3
-0.01
0.08
0.16
Syria_TellQarassa_Umayyad.SG
Israel_C.AG
Jordan_PPN.AG
Israel_Natufian.AG
Turkey_Marmara_Ilipinar_N.AG
NorthMacedonia_N.AG
Jordan_Baja_Late_PPNB.AG
Turkey_Marmara_Illipinar_N.AG
Croatia_N_Starcevo.AG
Jordan_Baja_PPNB.AG
BedouinB.DG
Turkey_Hatay_TellKurdu_Chalcolithic.AG
Greece_Lazarides_EBA.AG
Kenya_HyraxHill_PN.AG
Egypt_ThirdIntermediatePeriod.AG
Kenya_LukenyaHill_PN.AG
Serbia_VincaBelo_Starcevo_EN.SG
Jordan_EBA.AG
Hungary_N_Koros.AG
Croatia_MN.AG
Hungary_MN_LBK.SG
Congo_NgongoMbata_Protohistoric.AG
Tanzania_Lindi_Swahili.AG
South_Africa_2000BP.SG
Uganda_Munsa_LIA.AG
Congo_Kindoki_Protohistoric.AG
YRI_o.SG
BantuSA_Herero.DG
ESN.DG
YRI.DG
BantuHerero.DG

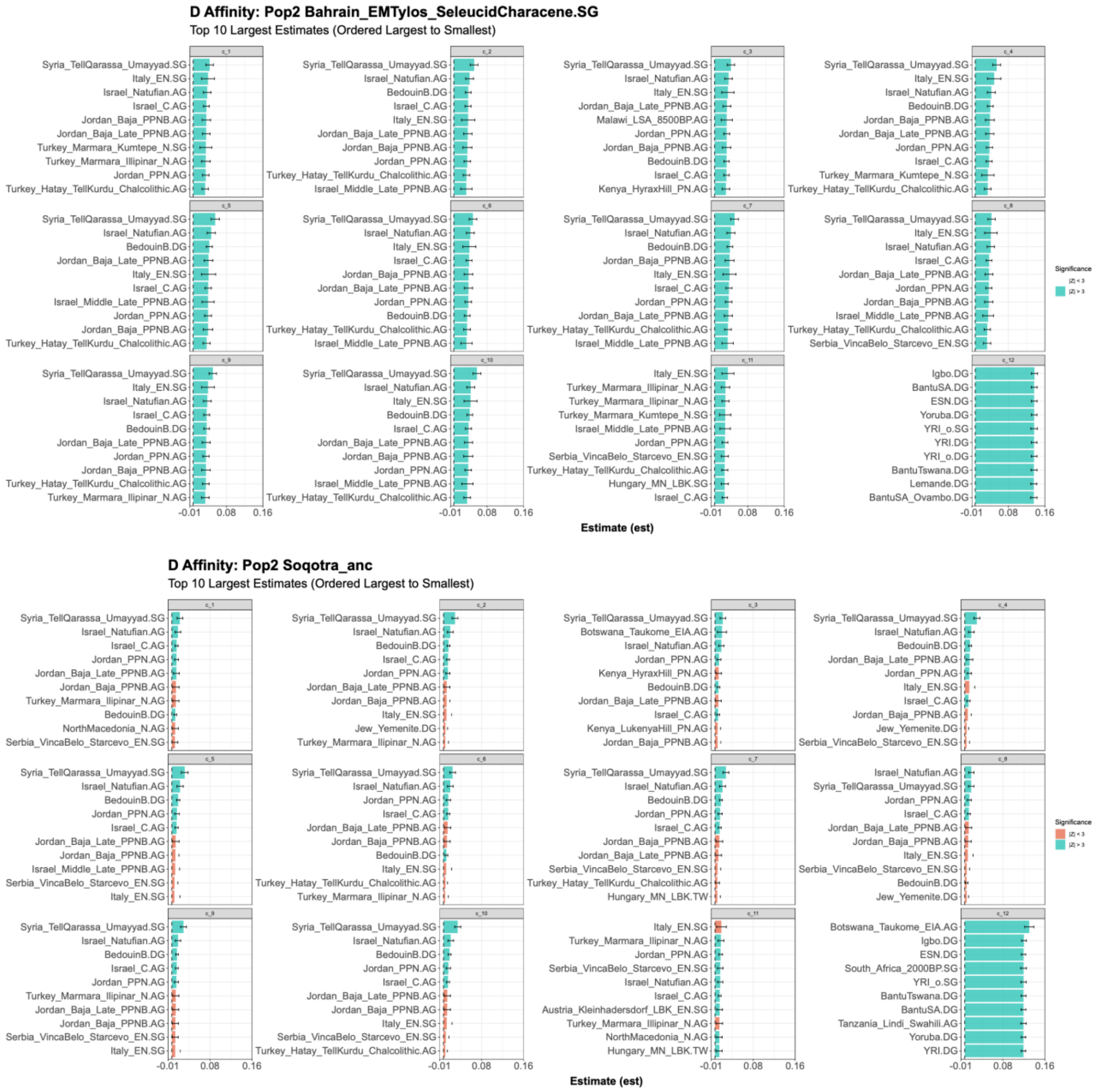


**Figure S5: Affinity D-statistics to test whether Saudi populations share more drift with ancient Arabian groups relative to diverse Eurasian and African populations.** We computed the D-statistics of the form D(WorldPop, ancient Arabian; SaudiCluster, Karitiana.DG). A significantly positive D-statistics with Z-score > 3 would suggest the Saudi populations share more drift with diverse Eurasian and/or African WorldPop relative to with ancient Arabians. Teal bars indicate statistically significant affinity estimates (|Z| > 3), while coral bars represent non-significant results (|Z| < 3). Error bars indicate +/- 1 standard error. These analyses revealed Saudi clusters 1:10 are highly consistent across all four ancient-Arabian-sources in revealing increased genetic affinity of Saudi clusters toward Levantine sources (frequently including Syria_TellQarassa_Umayyad.SG as the top hit along with Israel_Natufian.AG, Israel_C.AG, and a set of Levantine Neolithic/EBA, Jordan_PPN.AG, Jordan_Baja_PPNB.AG, Jordan_Baja_Late_PPNB.AG, Jordan_EBA.AG, BedouinB.DG) relative to ancient Bahrain (positive D-statistics with Z-score >3). The affinity D-statistics also reveal differences between cluster 11, cluster 12 from the remainder of the Saudi clusters. Saudi cluster 12 exhibits the most pronounced divergence from the baseline across all ancient-source selections, with its top-10 D-statistic rankings dominated by Sub-Saharan African and African-adjacent ancient or historic populations rather than Levantine groups. Saudi cluster 11 deviates more subtly but consistently by shifting its strongest affinities away from the Levant-only baseline toward a mixed Anatolia + European Neolithic/Mediterranean set. More subtly, Saudi cluster 3 appears to both possess the Levantine-related ancestry component, in addition to East African pastoralist/forager proxies as included in the top-10 populations.

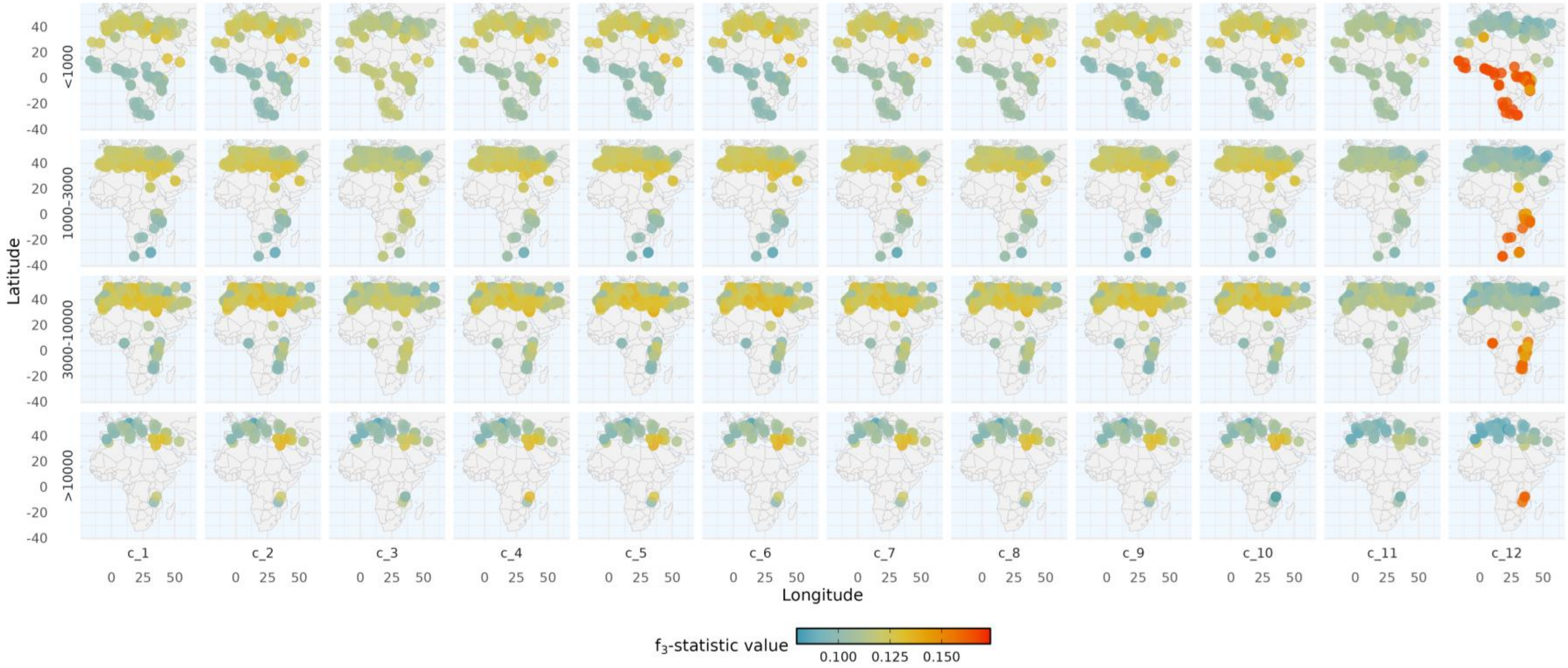


**Figure S6: Spatiotemporal mapping of shared genetic drift between modern Saudi clusters and diverse global populations.** Geospatial heatmaps display outgroup f3-statistics of the form f3(Han.DG; SaudiCluster, WorldPop), computed using *qp3pop* to quantify genetic affinities. The panels illustrate the shared genetic drift between distinct present-day Saudi genetic clusters (columns, c_1 to c_12) and 1,698 comparative Eurasian and African populations. The analysis is stratified across four distinct temporal periods based on the mean sample age of the test populations (rows: <1,000; 1,000–3,000; 3,000–10,000; and >10,000 years BP). Point colors correspond to the magnitude of the f3-statistic, with warmer colors indicating higher shared genetic drift relative to the Han.DG outgroup. This geospatial modeling highlights distinct regional affinity signatures and temporal shifts across the subpopulations, notably illustrating the pronounced and deep African affinity unique to cluster c_12.

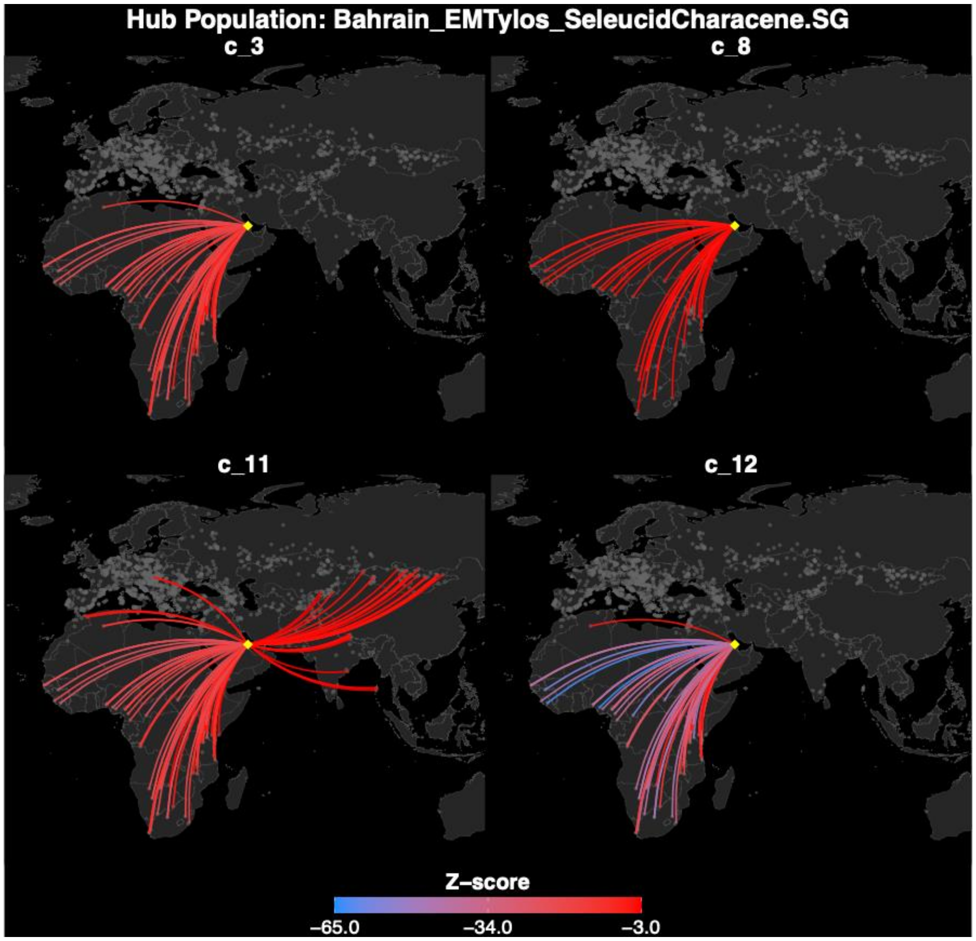
Hub Population: Bahrain_EMTylos_SeleucidCharacene.SG
c_3
c_8
c_11
c_12
Z–score
–65.0
–34.0
–3.0

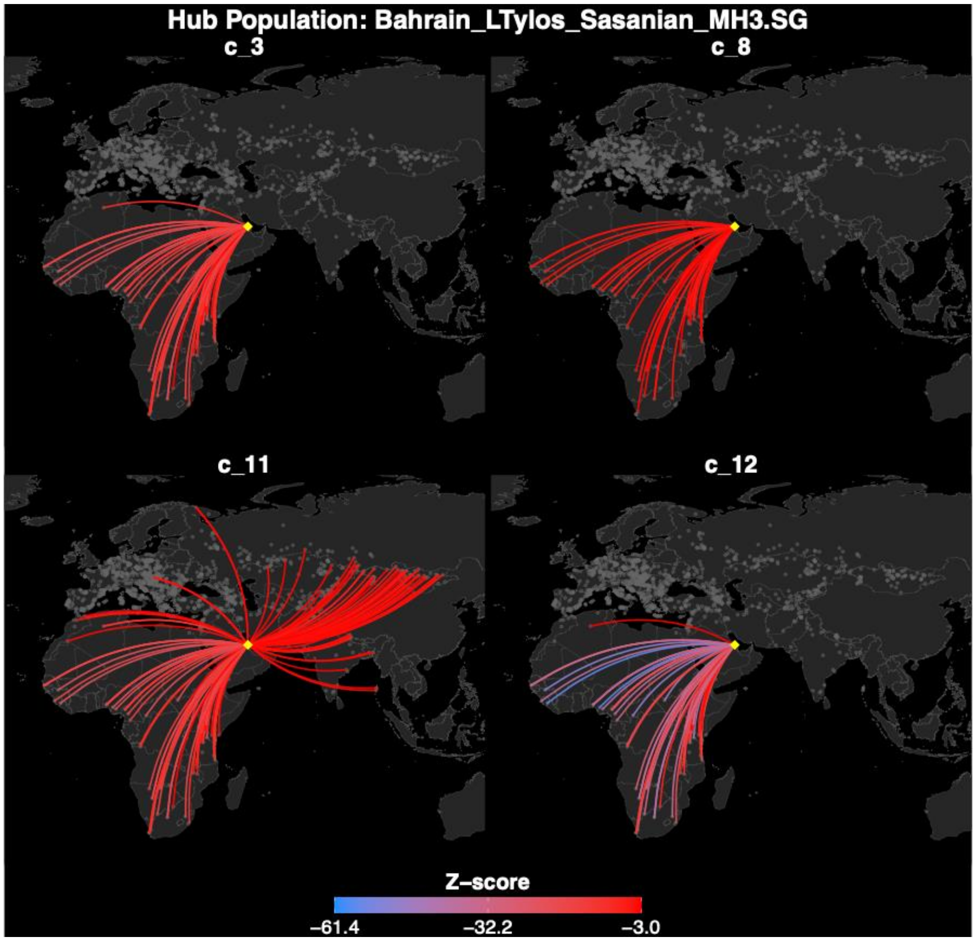
Hub Population: Bahrain_LTylos_Sasanian_MH3.SG
c_3
c_8
c_11
c_12
Z–score
–61.4
–32.2
–3.0

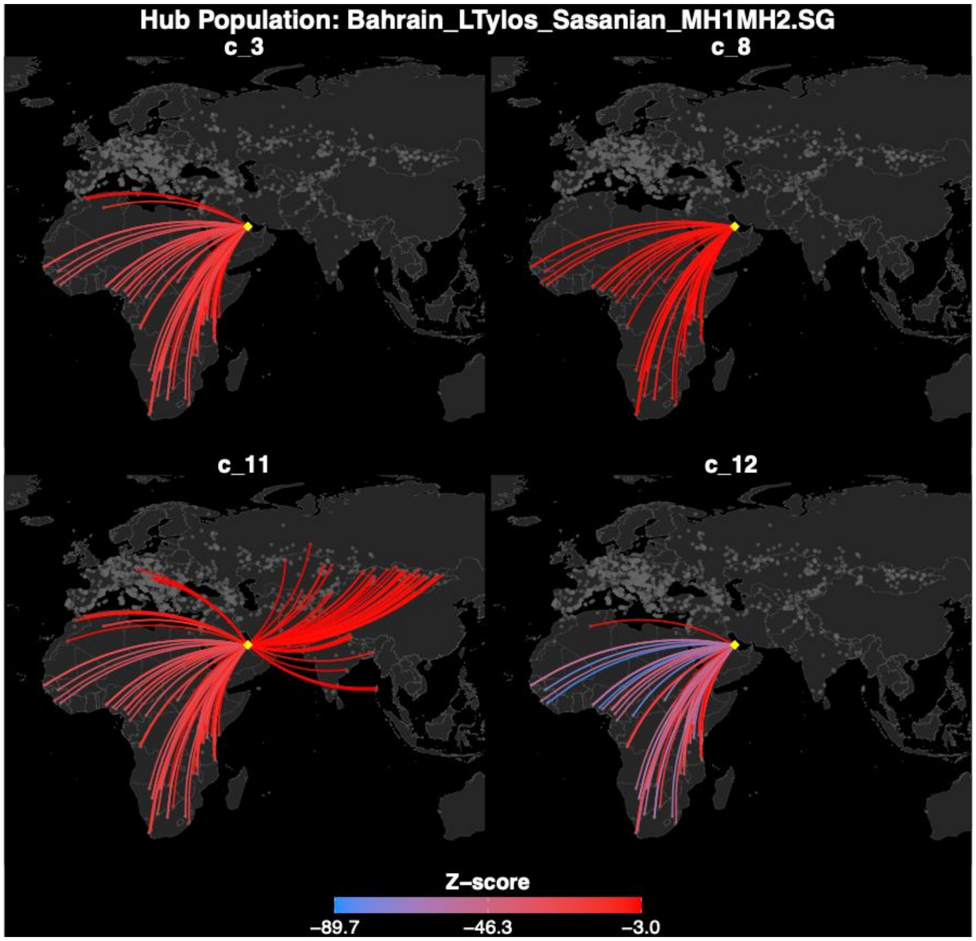

Hub Population: Bahrain_LTylos_Sasanian_MH1MH2.SG
c_3
c_8
c_11
c_12
Z-score
–89.7
–46.3
–3.0

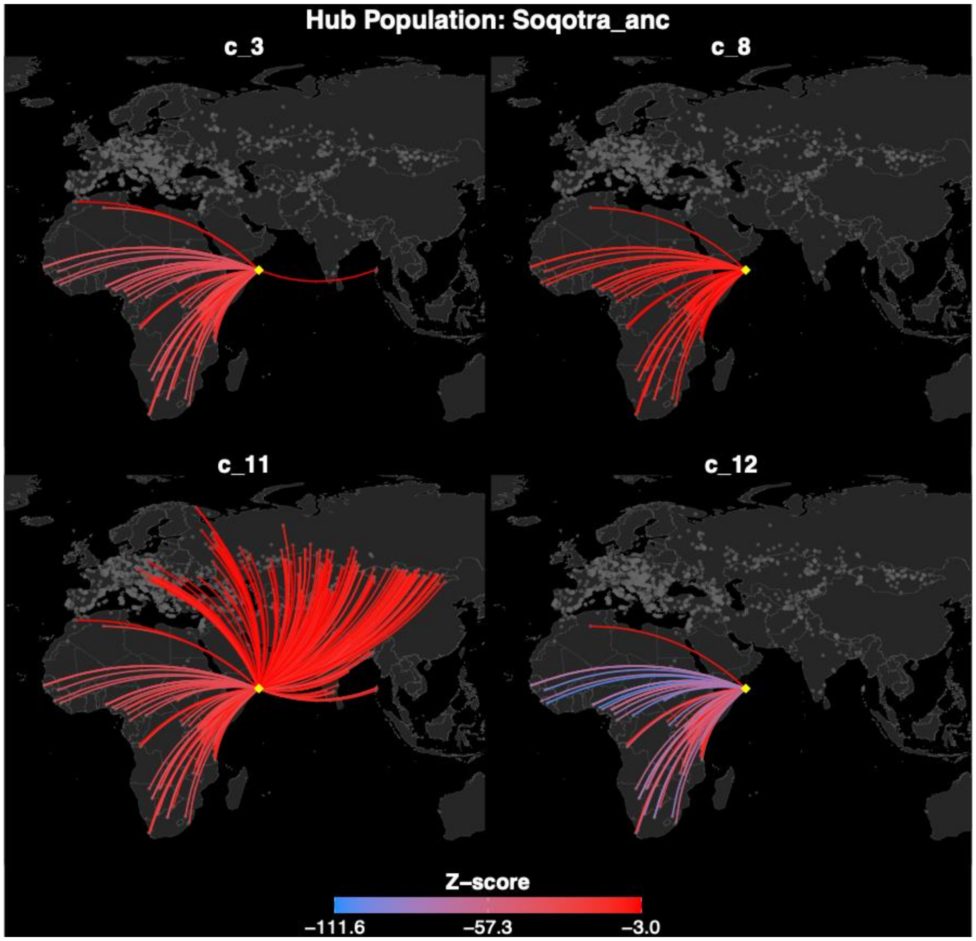


**Figure S7. Admixture f3 elucidate connections to Africa and Eastern Europe/Steppe.** We formally tested whether Saudi populations represent admixtures between ancient Arabian and global populations using admixture $f_3$-statistics of form $f_3$(SaudiCluster; WorldPop, ancient Arabian), where we fixed one of the four ancient Arabian populations and cycled through ancient and present-day global populations as possible second sources. As such, significantly negative values (Z < -3) indicate Saudi clusters derive from lineages related to the tested ancient Arabian and WorldPop population sources. Interestingly, of all $f_3$-statistic tests only four of the twelve Saudi clusters (3, 8, 11, and 12) yielded a significant admixture $f_3$-statistic. We visualized the patterns of unique and shared admixture $f_3$-statistic estimates between the Saudi clusters across each of the four ancient Arabian populations through "flight-path" maps. Here for each ancient Arabian source (yellow diamond), we visualized the significant (Z < -3) connections between them and the WorldPop as a line, with line color scaling with the magnitude of the Z-score. This allows us to visualize which pairs of ancient Arabian and Ancient-Global populations are shared signatures of admixture in the four Saudi clusters and which represent unique admixture sources.

**A**

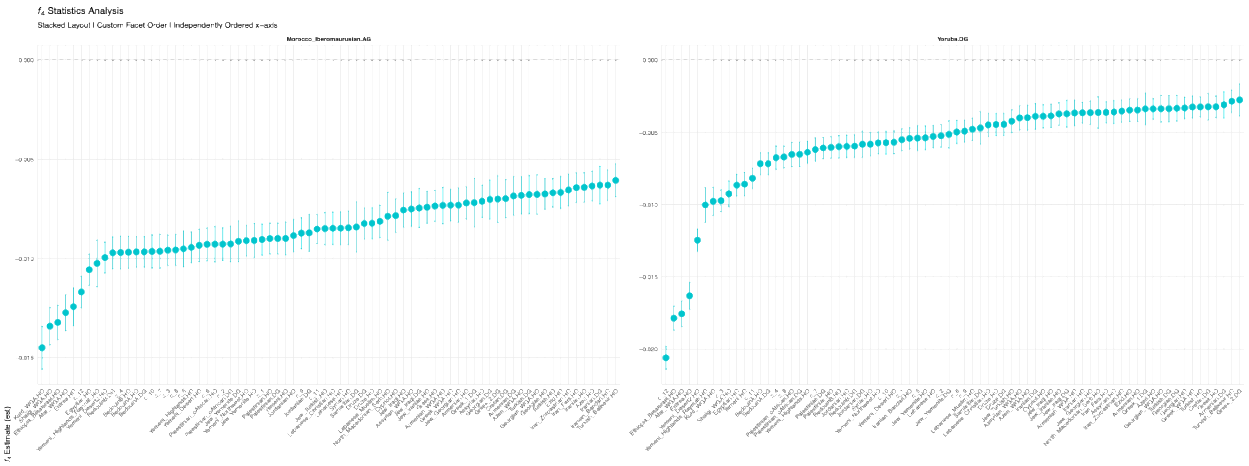


**B**

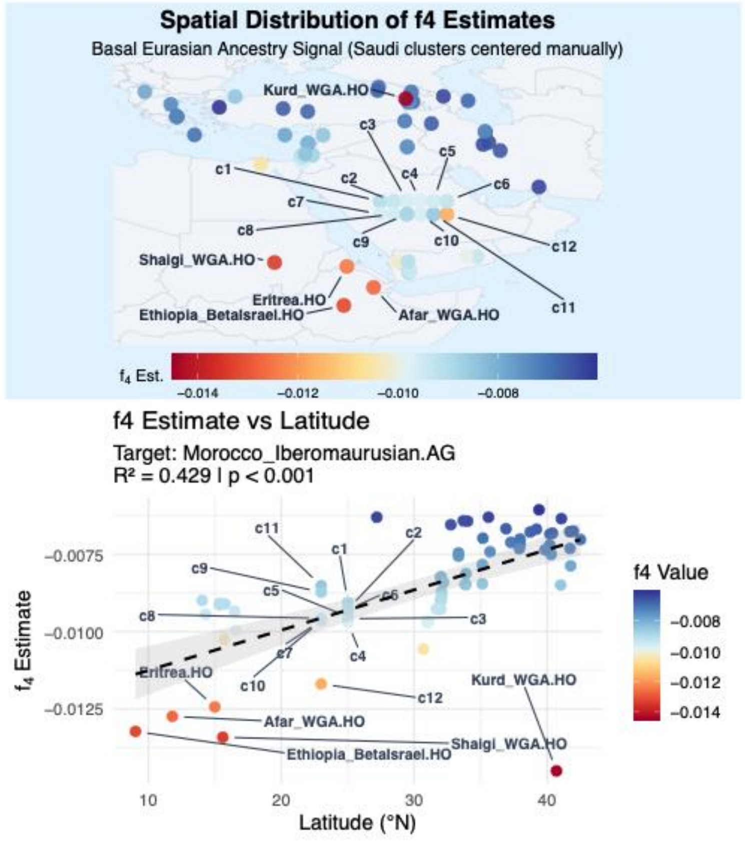


**Figure S8. Estimates of relative Basal Eurasian ancestry in Saudi clusters and surrounding modern populations.** (A) We used the $f_4$-statistic of form $f_4$(Saudi cluster, Han.DG; Ust-Ishim, Outgroups) to estimate the relative amount of Basal Eurasian ancestry (i.e. the drift basal to the shared drift between Han.DG and Ust-Ishim, a 45 ky sample from western Siberia) across the Saudi cluster cohort. Strongly negative $f_4$ values of this form is consistent with elevated shifted drift with Basal Eurasian ancestry. However, recent African admixture could confound this signal, and different African aDNA samples are likely to represent Basal Eurasian ancestry to different degrees, thereby prompting us to calculate $f_4$ using either Yoruba.DG and Morocco_Iberomaurusian.AG as outgroups. (B) Overall, we observe a latitudinal cline where Basal Eurasian proportions decrease in the northern Middle East, and a gradient within Saudi. Note all Saudi clusters were assigned the same latitude. One outlier is the Kurd sample, where they showed significant Basal Eurasian proportions considering their more Northern location.

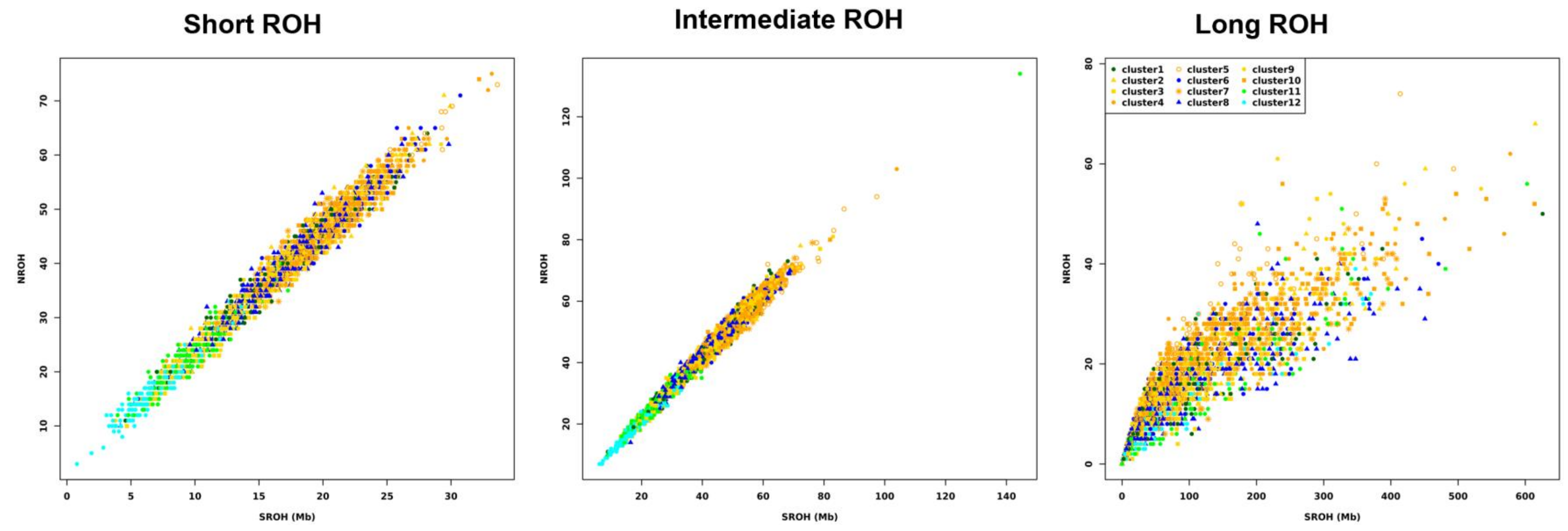


**Figure S9. Total length and number of ROH for different size classes.** Each individual is represented in the plot, labeled and colored based on their inferred cluster membership. For both the short and intermediate ROHs, there are clear linear relationships between NROH and SROH. In contrast, for the long ROHs, as SROH increase per individual genome, the NROHs are not increasing at the similar linear pattern as observed for short and intermediate ROHs. That is, for individuals with greater SROH due to the long ROHs, they do not have proportionally greater NROH compared to those with less SROH, suggesting that the contributions of SROHs are driven by fewer but longer ROHs in this length class due to recent consanguinity.

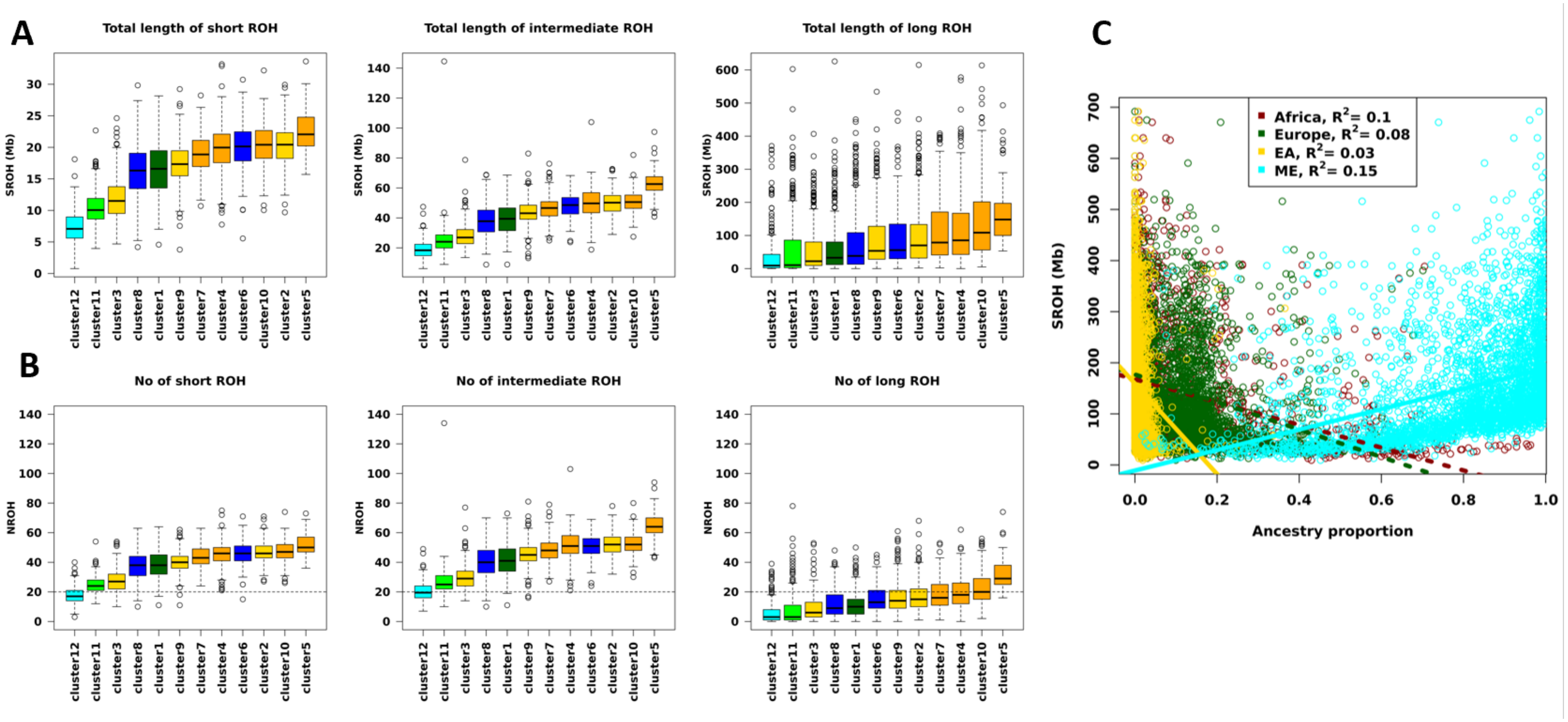


**Figure S10. Runs of homozygosity in Saudi Arabians.** The (A) total sum length of ROH and (B) Number of ROH are shown for each Saudi cluster, stratified by size classes. (C) The total sum of length of ROH per individual, across all ROHs, stratified by ancestry proportions. ROH – Runs of homozygosity, ME -Middle Eastern, EA – East Asia.

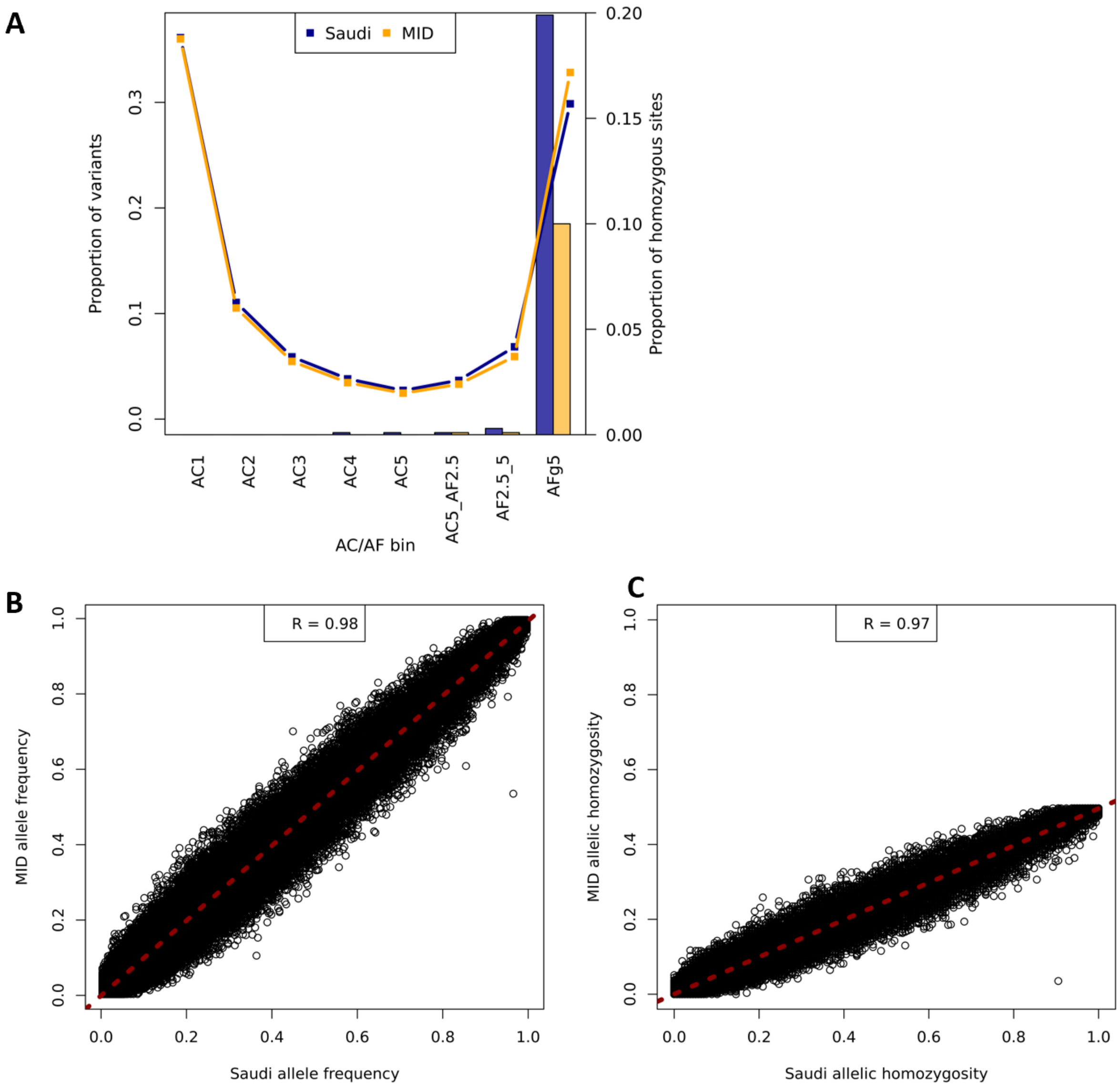


**Figure S11. Distribution of alternative allele frequency and homozygosity in Saudi vs gnomAD-MID.** (A) Genome-wide alternative allele frequency spectrum of Saudi and gnomAD-MID, and proportion of homozygous alternative allele per frequency bin. The sample size of Saudi is based on downsampling to gnomAD-MID sample size, n = 158. (B) Alternative allele frequency of Saudi vs. gnomAD-MID on shared segregating sites. (C) Homozygosity of alternative allele frequency of Saudi vs gnomAD-MID on shared segregating sites. The gnomAD-MID samples were based on 158 individuals with WGS from gnomAD v3 and the Saudi dataset was downsampled to 158 same sample for these comparisons. AC and AF refer to allele count and allele frequency, respectively. AFg5 refers to allele frequency greater than 5%. For better presentation, both (B) and (C) plots were based on randomly sampled subsets of 100,000 markers. MID – refers gnomAD-MID.

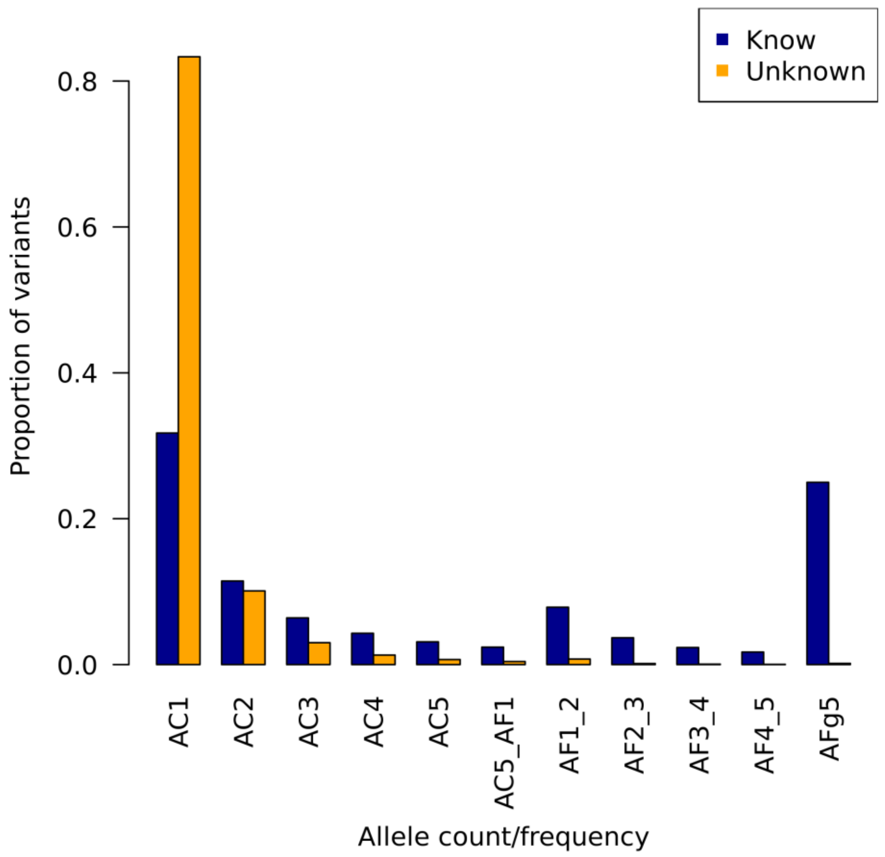


**Figure S12**. **Genome-wide distribution of minor allele frequency in Saudi population.** The 'known' (n = 23,029,031) variants are those that were found in gnomAD V4.1 and the 'unknown' (n = 2,459,950) are those that were not found in gnomAD. AC 1 to 5 refers to variants with allele count of 1 to 5, respectively, in our dataset. AF refers to allele frequency, and the numbers 1 to 5 next to AF represent percentages, e.g. AF1_2 refers to variants with allele frequencies between 1% and 2%. AFg5 refers to allele frequency greater than 5%.

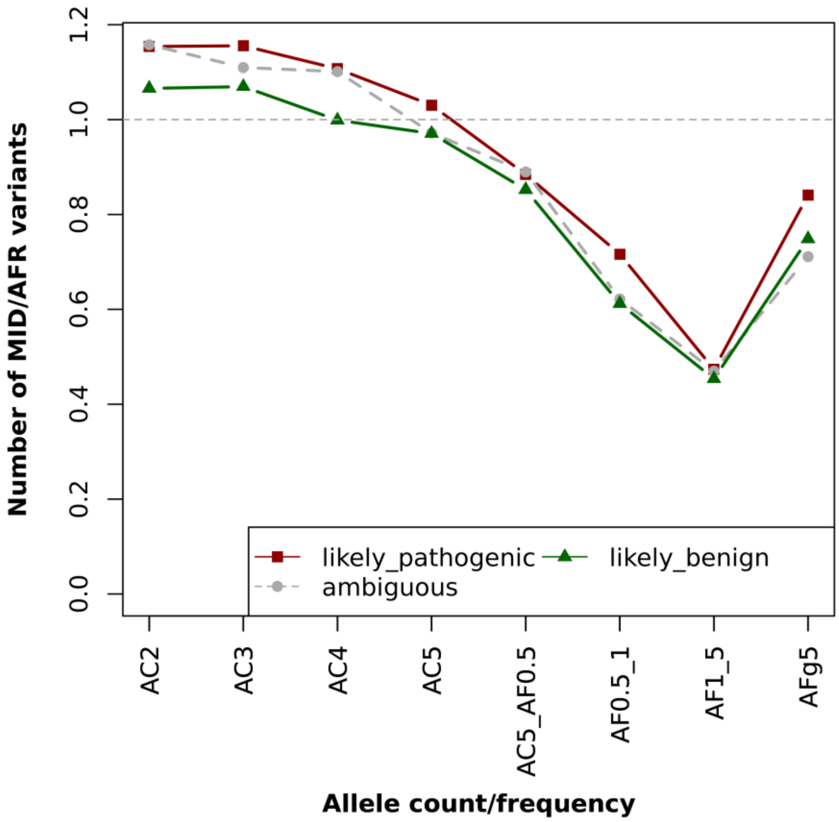


**Figure S13. Distribution of minor allele frequency between gnomAD-MID and gnomAD-AFR populations with variants annotated by AlphaMissense.** This analysis took advantage of the larger sample sizes for gnomAD-MID exome available, as well as the benefit of both comparison populations being processed through the same gnomAD pipeline. The sample size of gnomAD-AFR is based on downsampling to gnomAD-MID sample size, n = 2,596. AC and AF refer to allele count and allele frequency, respectively. AFg5 refers to allele frequency greater than 5%. MID and AFR refer to gnomAD-MID and gnomAD-AFR samples, respectively. ** denotes frequency bins with significant difference between the most deleterious (red) and most neutral (green) through bootstrapping at $p < 0.01$.

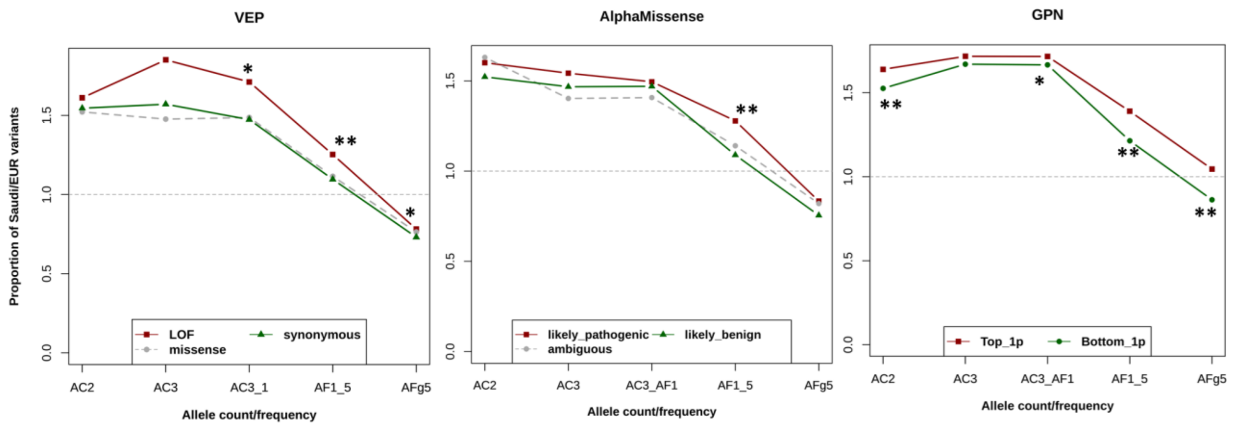


**Figure S14. Distribution of minor allele frequency between Saudi and gnomAD European populations.** The enrichment of deleterious alleles, compared to the benign alleles, appears to be more attenuated in this comparison than when Saudi was compared to the gnomAD-AFR population. The less significant finding when comparing Saudi to gnomAD-EUR is probably because Europeans also showed proportionally more deleterious than neutral alleles across all frequency bins, as previously reported (1, 2) and replicated here (Figure S15). AC and AF refer to allele count and allele frequency, respectively. AFg5 refers to allele frequency greater than 5%. Top_1p refers to variants with the top 1% of GPN scores (more deleterious) and Bottom_1p refers to variants with the bottom 1% of GPN scores (more neutral). EUR denotes the gnomAD-EUR samples. The sample size of gnomAD-EUR is based on downsampling to Saudi sample size, n = 302. LOF refers to Loss of function. ** and * denote frequency bins with significant difference between the most deleterious (red) and most neutral (green) through bootstrapping at $p < 0.01$ and $< 0.05$, respectively.

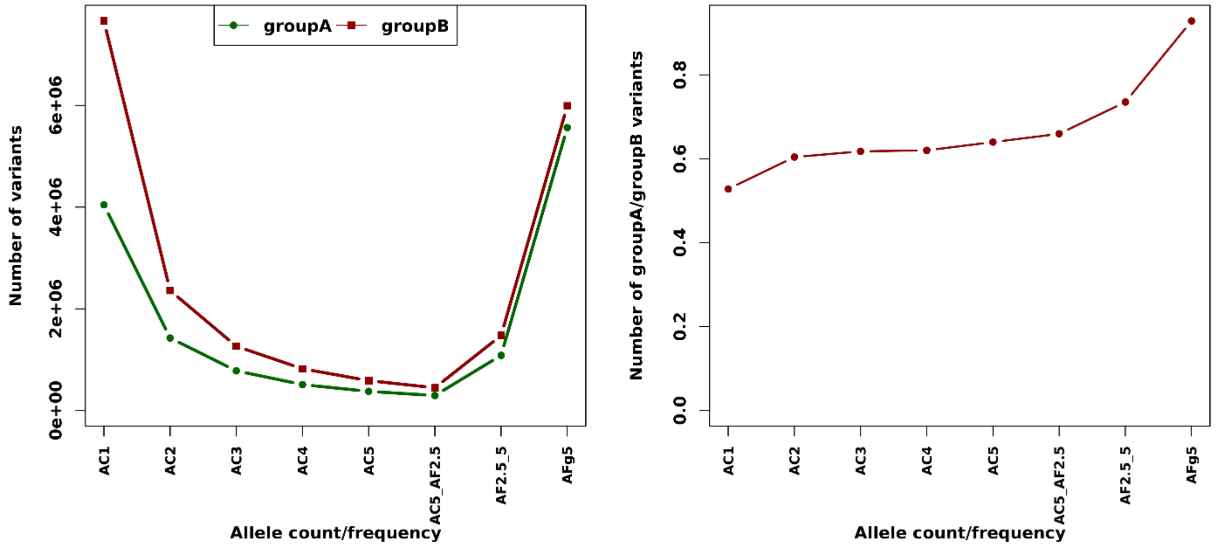


**Figure S15**. **Genome-wide distribution of allele frequencies in Saudi clusters.** (A) comparison of allele frequency between cluster groupA and groupB, (B) ratio of number of variants of groupA to groupB clusters. The sample size of cluster groupB is based on downsampling to groupA sample size, n = 124. AC and AF refer to allele count and allele frequency, respectively. AFg5 refers to allele frequency greater than 5%.

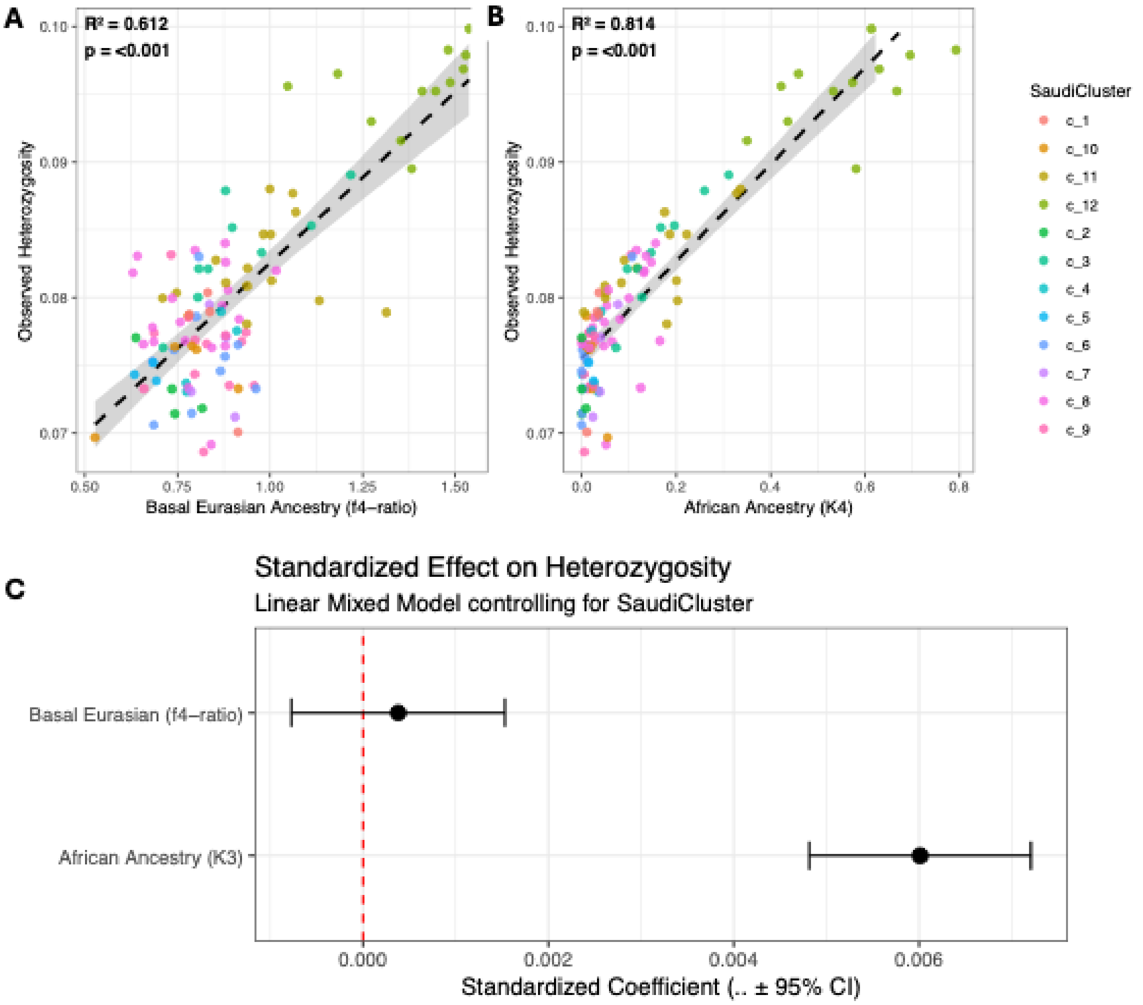


**D**

Effect of Ancestry on Heterozygosity

| Predictor | Std. Beta | Std. Error | t-value | p-value (LMM) | VIF | Partial R2 (LMM) | Partial r (adj) | p-value (Partial r) |
|---|---|---|---|---|---|---|---|---|
| Basal Eurasian (f4-ratio) | 4e-04 | 6e-04 | 0.647 | 0.519 | 3.62 | 0.0002 | -0.0010 | 0.992 |
| African Ancestry (K4) | 6e-03 | 6e-04 | 9.851 | <0.001 | 3.62 | 0.6984 | 0.4865 | <0.001 |

LMM Marginal R²: 0.805 | Conditional R²: 0.816

**E**

Nested Model Comparisons

Likelihood Ratio Tests evaluating model improvement when adding ancestry predictors

| Base Model | Added Predictor | AIC (Base Model) | AIC (Full Model) | Chi-Square | p-value |
|---|---|---|---|---|---|
| Basal Eurasian Only | African Ancestry (K4) | -840.70 | -890.86 | 52.160 | <0.001 |
| African Ancestry Only | Basal Eurasian (f4-ratio) | -892.41 | -890.86 | 0.455 | 0.5 |

A lower AIC indicates a better fitting model. The p-value tests if the Full Model (Basal + African) is significantly better than the Base Model.

**Figure S16. Evaluating the effects of Basal Eurasian and African-like ancestries on genomic diversity in Saudi individuals.** Univariate relationships between ancestry and observed heterozygosity. Scatter plots with dashed linear regression lines showing the correlation between an individual's observed heterozygosity and (A) Basal Eurasian ancestry (estimated via an f4-ratio utilizing a Morocco Iberomaurusian reference; **Methods**), and (B) African-like ancestry, estimated via unsupervised ADMIXTURE (K = 4; **Figure 1C**). (C) Standardized effect sizes from Linear Mixed Modeling (LMM). A forest plot displaying the standardized coefficients and 95% confidence intervals from a LMM model that regresses heterozygosity against both scaled ancestry predictors simultaneously, controlling for subpopulation structure by treating the Saudi cluster as a random intercept. After controlling for African ancestry, the effect of Basal Eurasian ancestry is indistinguishable from zero. (D) LMM summary statistics, variance partitioning, and partial correlations. Detailed metrics for the model shown in (C). Multicollinearity diagnostics confirm the model is stable (VIF = 3.62, below the critical threshold of 5). Partial (derived via nested model subtraction) and within-cluster partial correlations (derived via Frisch-Waugh-Lovell residualization) demonstrate that African ancestry uniquely explains ~69.8% of the variance in heterozygosity (Partial $R^2$ = 0.6984, Partial $r_{adj}$ = 0.4865, p<0.001). Conversely, Basal Eurasian ancestry explains negligible unique variance (Partial $R^2$ = 0.0002). Overall, the fixed ancestry effects explain 80.5% of total heterozygosity variance (Marginal $R^2$ = 0.805). (E) Nested model comparisons via Likelihood Ratio Tests. Analysis of Variance (ANOVA) evaluating whether adding an ancestry component significantly improves predictive fit. Adding African ancestry to a Basal Eurasian-only base model yields a significant improvement in fit (AIC drops from -840.70 to -890.86; $X^2$ = 52.16, p<0.001). Conversely, adding Basal Eurasian ancestry to an African-only base model yields no significant improvement (*p*=0.50), confirming that Basal Eurasian ancestry provides no independent explanatory power for modeling genomic diversity in this cohort.

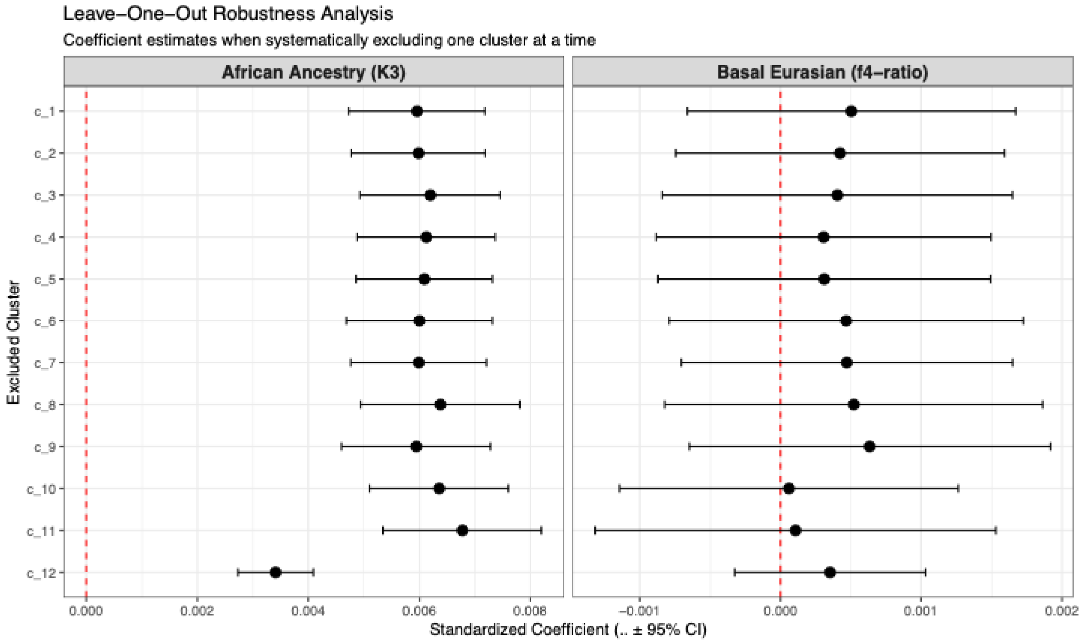


### Leave-One-Out (LOO) Model Robustness

Assessing if any single cluster drives the Ancestry-Heterozygosity relationship

| Excluded | Basal β | Basal t-val | K3 β | K3 t-val | Model R² | K3 Shift (Z) | Outlier? |
|---|---|---|---|---|---|---|---|
| c_1 | 0.0005 | 0.85 | 0.0060 | 9.52 | 0.8097 | 0.0284 | No |
| c_2 | 0.0004 | 0.71 | 0.0060 | 9.73 | 0.8061 | 0.0592 | No |
| c_3 | 0.0004 | 0.64 | 0.0062 | 9.63 | 0.8124 | 0.3161 | No |
| c_4 | 0.0003 | 0.51 | 0.0061 | 9.69 | 0.8045 | 0.2295 | No |
| c_5 | 0.0003 | 0.52 | 0.0061 | 9.75 | 0.8019 | 0.1870 | No |
| c_6 | 0.0005 | 0.73 | 0.0060 | 8.96 | 0.8020 | 0.0794 | No |
| c_7 | 0.0005 | 0.78 | 0.0060 | 9.64 | 0.8090 | 0.0655 | No |
| c_8 | 0.0005 | 0.76 | 0.0064 | 8.73 | 0.8287 | 0.5352 | No |
| c_9 | 0.0006 | 0.97 | 0.0059 | 8.69 | 0.8080 | 0.0125 | No |
| c_10 | 0.0001 | 0.10 | 0.0064 | 9.97 | 0.8105 | 0.5094 | No |
| c_11 | 0.0001 | 0.15 | 0.0068 | 9.31 | 0.8273 | 1.0146 | No |
| c_12 | 0.0004 | 1.02 | 0.0034 | 9.83 | 0.6069 | 3.0368 | Yes |

K3 Shift (Z) > 2 indicates that removing the cluster significantly alters the African Ancestry effect size relative to the LOO average.

**Figure S17. Sensitivity analysis of ancestry effects on genomic diversity via linear mixed models.** The linear mixed model analysis in **Figure S16** were repeated here, each time leaving out one Saudi cluster at a time, to test the robustness of the LMM coefficient estimates. The forest plots and corresponding table display the shift in standardized effect sizes (beta-coefficient) for both ancestry components when systematically excluding one cluster at a time. The analysis reveals that cluster c_12 acts as a highly influential outlier; its exclusion significantly alters the effect size of African-like ancestry on heterozygosity (K3 Shift Z > 2), indicating that the strength of this specific relationship is disproportionately driven by the c_12 subpopulation. Nevertheless, in all cases Basal Eurasian ancestry is not significantly associated with heterozygosity after accounting for African-like ancestry.

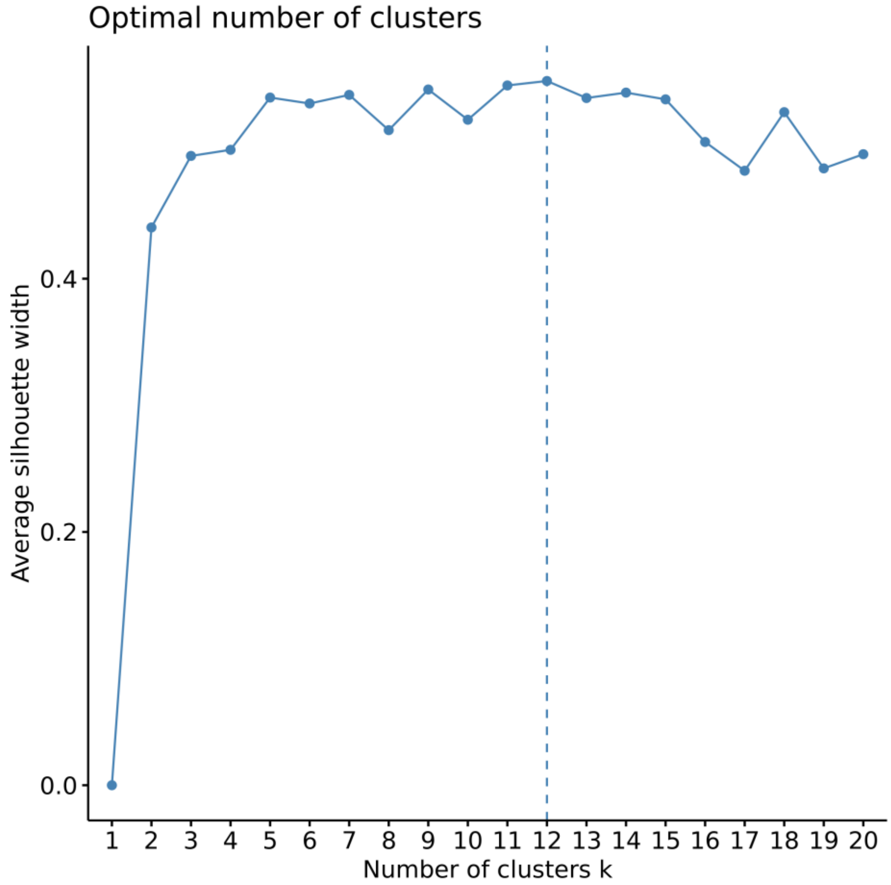


**Figure S18. Estimated optimal number of clusters using Average Silhouette Width.** The maximum value was achieved at k = 12, thought k = 5 or k = 9 could be equally sensible.

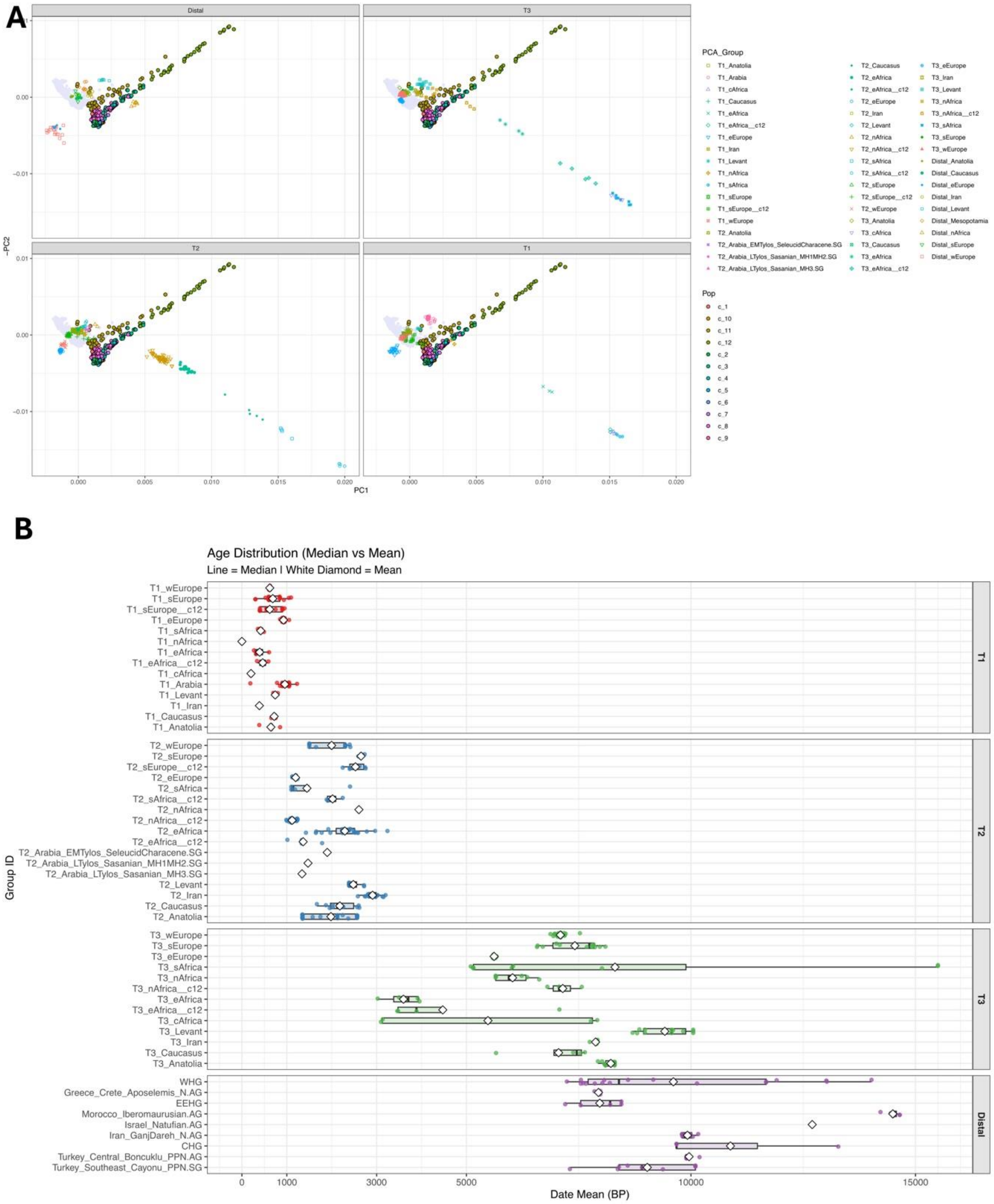


**Figure S19. Principal component analysis and temporal distribution of meta-population sources constructed for *qpAdm* modeling.** (A) Principal Component Analysis (PCA) projecting ancient genomes (HO AADR v.62.0) onto the genetic variation of 70 modern global populations and present-day Saudi clusters. To define meta-population sources for downstream *qpAdm*

analysis, populations within each temporal period (Distal, T4, T2, and T1) were merged based on their clustering in PCA space and corresponding high outgroup f3-statistic affinities. PCA was performed using *smartpca* (EIGENSOFT) with least-squares projection (lsqproject: YES). (B) Age distribution (in years BP) of the resulting meta-population groups utilized as sources, stratified by temporal period (T1, T2, T3, and Distal). For each meta-population, the boxplot illustrates the range of sample dates, with vertical lines indicating the median date and white diamonds denoting the mean date. Full population compositions corresponding to these meta-population groupings are detailed in **Table S14**.
