## Supplementary Tables for "Patterns of population structure and genetic variation within the Saudi Arabian population"

### Table S1. Distribution of Saudi samples by inferred sub-clusters and tribal region

| | Central | West | North | South | East | North_ Central | Total | WGS |
|---|---|---|---|---|---|---|---|---|
| **cluster1** | 28 [8] | 13 [1] | **269** [22] | 11 [5] | 0 [0] | 0 [0] | **321 [36]** | 20 |
| **cluster2** | **164** [17] | 94 [6] | 0 [0] | 19 [9] | 0 [0] | 0 [0] | **277 [32]** | 16 |
| **cluster3** | **293** [39] | 7 [13] | 0 [4] | 1 [0] | 0 [1] | 0 [1] | **301 [58]** | 33 |
| **cluster4** | 3 [3] | **304** [30] | 0 [0] | 0 [3] | 0 [1] | 0 [0] | **307 [37]** | 9 |
| **cluster5** | 0 [0] | **57** [13] | 0 [2] | 0 [3] | 0 [1] | 0 [0] | **57 [19]** | 7 |
| **cluster6** | 19 [4] | 2 [1] | 0 [0] | **221** [17] | 0 [2] | 0 [0] | **242 [24]** | 19 |
| **cluster7** | 1 [1] | **242** [61] | 109 [15] | 1 [5] | 0 [3] | 0 [0] | **353 [85]** | 8 |
| **cluster8** | 10 [13] | 33 [19] | 0 [3] | **338** [44] | 0 [6] | 0 [0] | **381 [85]** | 59 |
| **cluster9** | **329** [56] | 15 [14] | 3 [11] | 6 [6] | 3 [8] | 0 [0] | **356 [95]** | 48 |
| **cluster10** | 1 [0] | **302** [18] | 0 [0] | 0 [0] | 0 [0] | 0 [0] | **303 [18]** | 17 |
| **cluster11** | 82 [19] | 80 [19] | 39 [8] | 31 [13] | 10 [9] | 0 [0] | **242 [68]** | 36 |
| **cluster12** | 92 [19] | 56 [13] | 0 [3] | 64 [16] | 0 [4] | 0 [0] | **212 [55]** | 30 |
| **Total** | **1, 022 [179]** | **1, 205 [208]** | **420 [68]** | **692 [121]** | **13 [35]** | **0 [1]** | **3, 352 [612]** | **302** |

For each column other than WGS, number of individuals based on HARE-imputed label is followed by number of self-reported labels in brackets. None of the WGS samples had tribal information nor were tribal information imputed, but their distribution in each cluster is shown.

**Table S2. Average admixture proportions based on K=4**

| | Presumed ancestry | | | |
|---|---|---|---|---|
| | **EA-like** | **AFR-like** | **EUR-like** | **ME-like** |
| **Saudi clusters** | | | | |
| cluster1 | 0.010 | 0.037 | **0.165** | **0.788** |
| cluster2 | 0.002 | 0.007 | 0.020 | **0.971** |
| cluster3 | 0.029 | **0.197** | **0.143** | **0.631** |
| cluster4 | 0.003 | 0.027 | 0.013 | **0.957** |
| cluster5 | 0.002 | 0.010 | 0.006 | **0.983** |
| cluster6 | 0.002 | 0.006 | 0.078 | **0.914** |
| cluster7 | 0.003 | 0.037 | 0.038 | **0.922** |
| cluster8 | 0.009 | 0.083 | **0.161** | **0.747** |
| cluster9 | 0.013 | 0.013 | **0.141** | **0.833** |
| cluster10 | 0.003 | 0.015 | 0.002 | **0.980** |
| cluster11 | 0.048 | **0.144** | **0.364** | **0.444** |
| cluster12 | 0.024 | **0.549** | 0.074 | **0.354** |
| **HGDP Middle Easterners** | | | | |
| BedouinA | 0.014 | 0.111 | **0.396** | **0.479** |
| BedouinB | 0.003 | 0.078 | 0.089 | **0.830** |
| Druze | 0.013 | 0.028 | 0.590 | 0.370 |
| Palestinian | 0.010 | 0.088 | 0.473 | 0.429 |
| Mozabite | 0.000 | 0.289 | 0.397 | 0.314 |

We labelled the ancestries by the HGDP population or group of individuals with dominating or highest admixture proportions. EA – East Asian, ME - Middle Eastern, AFR - African, EUR – European. Bedouin, particularly BedouinB, shows the highest amount of the ME-like ancestry, comparable to what we observe in the Saudi data.

**Table S3. Average admixture proportions based on K=9**

| | Presumed ancestry | | | | | | | | |
|---|---|---|---|---|---|---|---|---|---|
| | **EA** | **Ocean** | **AFR** | **ME-1** | **ME-2** | **ME-3** | **AMR** | **EUR** | **CSA** |
| **Saudi clusters** | | | | | | | | | |
| cluster1 | 0.003 | 0.003 | 0.032 | **0.178** | **0.575** | 0.105 | 0.002 | 0.032 | 0.070 |
| cluster2 | 0.001 | 0.002 | 0.005 | 0.154 | **0.381** | **0.452** | 0.001 | 0.002 | 0.003 |
| cluster3 | 0.009 | 0.006 | **0.189** | **0.362** | **0.225** | 0.115 | 0.003 | 0.010 | 0.082 |
| cluster4 | 0.002 | 0.003 | 0.023 | 0.042 | **0.150** | **0.771** | 0.001 | 0.004 | 0.006 |
| cluster5 | 0.001 | 0.000 | 0.006 | **0.329** | **0.159** | **0.503** | 0.000 | 0.001 | 0.001 |
| cluster6 | 0.002 | 0.003 | 0.005 | 0.020 | **0.896** | 0.059 | 0.002 | 0.002 | 0.010 |
| cluster7 | 0.002 | 0.003 | 0.033 | 0.036 | **0.569** | **0.344** | 0.002 | 0.004 | 0.008 |
| cluster8 | 0.004 | 0.005 | 0.078 | 0.021 | **0.729** | 0.061 | 0.002 | 0.028 | 0.073 |
| cluster9 | 0.001 | 0.003 | 0.009 | **0.805** | 0.098 | 0.069 | 0.001 | 0.003 | 0.010 |
| cluster10 | 0.002 | 0.003 | 0.009 | 0.01 | 0.051 | **0.918** | 0.001 | 0.002 | 0.004 |
| cluster11 | 0.015 | 0.007 | **0.133** | 0.089 | **0.356** | 0.045 | 0.003 | **0.128** | **0.223** |
| cluster12 | 0.015 | 0.006 | **0.543** | 0.047 | **0.292** | 0.042 | 0.001 | 0.012 | 0.041 |
| **HGDP Middle Easterners** | | | | | | | | | |
| Druze | 0.000 | 0.001 | 0.015 | 0.017 | 0.389 | 0.001 | 0.000 | 0.295 | 0.282 |
| BedouinA | 0.002 | 0.003 | 0.104 | 0.042 | 0.445 | 0.037 | 0.001 | 0.197 | 0.170 |
| BedouinB | 0.002 | 0.003 | 0.074 | **0.133** | 0.461 | **0.281** | 0.002 | 0.029 | 0.016 |
| Mozabite | 0.003 | 0.004 | 0.296 | 0.001 | 0.300 | 0.008 | 0.000 | 0.387 | 0.000 |
| Palestinian | 0.000 | 0.002 | 0.078 | 0.037 | 0.434 | 0.004 | 0.000 | 0.236 | 0.209 |
| **HGDP Europeans** | | | | | | | | | |
| Sardinian | 0.002 | 0.003 | 0.004 | 0.001 | **0.184** | 0.002 | 0.001 | 0.804 | 0.000 |
| Russian | 0.056 | 0.001 | 0.000 | 0.000 | 0.000 | 0.000 | 0.066 | 0.661 | 0.215 |
| Bergamo Italian | 0.001 | 0.002 | 0.000 | 0.013 | **0.114** | 0.007 | 0.018 | 0.703 | 0.142 |
| Tuscan | 0.000 | 0.002 | 0.002 | 0.020 | **0.154** | 0.004 | 0.012 | 0.633 | 0.172 |
| French | 0.000 | 0.001 | 0.000 | 0.008 | 0.032 | 0.014 | 0.033 | 0.751 | 0.160 |
| Basque | 0.007 | 0.007 | 0.001 | 0.01 | 0.011 | 0.013 | 0.026 | 0.897 | 0.028 |
| Adygei | 0.023 | 0.002 | 0.000 | 0.033 | **0.121** | 0.005 | 0.022 | 0.347 | **0.448** |
| Orcadian | 0.000 | 0.001 | 0.000 | 0.000 | 0.000 | 0.000 | 0.042 | 0.780 | **0.177** |

We labelled the ancestries by the HGDP population or group of individuals with dominating or highest admixture proportions. EA – East Asian, ME - Middle Eastern, AFR - African, EUR – European, AMR – American, Ocean – Oceania, CSA – Central & South Asian. K = 9 showed the lowest in-sample cross-validation error, and also introduced new ancestry components that distinguished the CSA-like ancestry from the EUR-like ancestry (see **Figure S3A**). At K = 9, we observed three ME-like ancestries (also see **Figure 1C,** bottom). One of the Middle Eastern-like ancestry (ME-2) that is dominant in several (> 10% in 10 out of the 12) clusters, particularly in clusters 6, 8, 1, and 7, is also found in Sardinians and other Italians & Adygei but completely missing in the Russians. The other Middle Eastern-like ancestries (ME-1 and ME-3) are found distributed in a subset of the clusters. In particular, the ME-3 ancestry is found in high proportions in clusters 10, 4, 5, and 2 (average proportions = 0.45 - 0.92), and was also found in HGDP-Bedouin (but absent from HGDP-Druze, HGDP-Mozabite and HGDP-Palestinian). This ancestry appears to be enriched in Qataris Bedouins and Saudi Arabians but not other Middle Eastern populations, and was suggested to reflect an indigenous Arab ancestry (1).

**Table S13. Distribution of SNPs by functional classes as annotated by VEP and AlphaMissense**

| VEP annotation | | | |
|---|---|---|---|
| **SNP functional class** | **Number of SNPs [%]** | **Variants found in gnomAD [%]** | **Previously unknown variants in gnomAD [%]** |
| **High impact / loss of function** | | | |
| Splice acceptor | 2,138 [0.0084] | 1,782 [0.008] | 356 [0.014] |
| Splice donor | 3,032 [0.012] | 2,559 [0.011] | 473 [0.019] |
| Stop gained | 2,456 [0.0096] | 1,954 [0.008] | 502 [0.020] |
| Stop lost | 744 [0.0029] | 298 [0.001] | 66 [0.003] |
| Start lost | 364 [0.0014] | 512 [0.002] | 232 [0.009] |
| **Total LOF** | 8,734 [0.034] | 7,105 [0.031] | 1,629 [0.066] |
| **Moderate impact** | | | |
| Missense | 139,135 [0.55] | 117,657 [0.51] | 21,478 [0.87] |
| **Low impact** | | | |
| Synonymous | 101,862 [0.40] | 92,290 [0.40] | 9,572 [0.39] |
| Coding sequence | 16 [0.00006] | 14 [0] | 2 [0] |
| Non coding exon transcript | 709,590 [2.78] | 639,176 [2.78] | 70,414 [2.86] |
| 5' UTR | 74,248 [0.29] | 63,239 [0.27] | 11,009 [0.45] |
| 3' UTR | 277,778 [1.09] | 249,101 [1.08] | 28,677 [1.17] |
| Intronic | 14,265,842 [55.97] | 12,904,477 [56.04] | 1,361,365 [55.34] |
| Upstream | 1,134,050 [4.45] | 1,028,529 [4.47] | 105,521 [4.29] |
| Downstream | 920,874 [3.61] | 838,032 [3.64] | 82,842 [3.37] |
| Intergenic | 7,856,812 [30.82] | 7,089,372 [30.78] | 767,440 [31.20] |
| **Total annotated SNPs** | **25,488,941** | **23,028,992 [90.3]** | **2,459,949 [9.7]** |
| **Total Saudi SNPs** | **25,488,981** | | |
| **AlphaMissense annotation** | | | |
| **SNP functional class** | **Number of SNPs [%]** | **Variants found in gnomAD [%]** | **Previously unknown variants in gnomAD [%]** |
| **Likely pathogenic** | 11,009 [9.17] | 7,265 [7.2] | 3,744 [19.63] |
| **Ambiguous** | 8,614 [7.18] | 6,484 [6.4] | 2,130 [11.17] |
| **Likely benign** | 100,389 [83.65] | 87,192 [86.4] | 13,197 [69.20] |
| **Total** | **120,012** | **100,941 [84.11]** | **19,071 [15.89]** |
| **GPN annotation** | | | |
| **SNP functional class** | **Number of SNPs** | | |
| **Top 1 percent variants** | 97,906 | | |
| **Bottom 1 percent variants** | 522,490 | | |

Percentages are given in brackets after the variant counts for each category. Top 1 percent variants are those with top 1% of GPN scores (more deleterious) and bottom 1% refers to variants with the bottom 1% of GPN scores (more neutral).
